# The NimB2 opsonin promotes *S. aureus* recognition by macrophages in *Drosophila melanogaster*

**DOI:** 10.64898/2026.08.28.747790

**Authors:** Prince Kumar Sah, Asya Dolgikh, Fanny Schüpfer, Jean Philippe Boquete, Samuel Rommelaere, Sérgio R Filipe, Bruno Lemaitre

## Abstract

Phagocytosis is a conserved effector process of innate immunity that enables specialized immune cells to detect, engulf, and degrade invading microbes as well as dying cells and cellular debris. Soluble opsonins enhance this process by coating target surfaces and promoting their recognition and uptake by phagocytic cells. In *Drosophila*, many phagocytic receptors of the Nimrod family have been characterized, but opsonins remain comparatively poorly understood. Here, we identified the secreted Nimrod protein NimB2 as an insect opsonin that specifically promotes the clearance of *Staphylococcus aureus* bacteria. NimB2 is produced by the fat body, secreted into the hemolymph, and required for resistance to *S. aureus* infection. *NimB2* null mutants showed reduced phagocytosis of *S. aureus* while maintaining Toll-dependent antimicrobial peptide expression. NimB2 promotes *S. aureus* binding to plasmatocytes (*Drosophila* macrophages). Using binding assays and bacterial cell wall mutants, we found that NimB2 recognizes a lipoteichoic acid (LTA)–dependent determinant on the *S. aureus* surface that allows the coating of the bacterial envelope. We further show that this coating enables efficient recognition by hemocytes via the phagocytic receptor Eater, which is necessary for NimB2-dependent binding. Moreover, Eater overexpression enhances NimB2-mediated association of *S. aureus* with hemocytes. Together, our results establish NimB2 as an insect opsonin that links a specific bacterial ligand exposed on the cell envelope to a phagocytic receptor.

## Introduction

*Drosophila melanogaster* has emerged as a powerful system to dissect innate immunity mechanisms *in vivo*. The fly immune system relies on two arms that act in a coordinated manner to restrict microbial infection: a humoral response and a cellular response (Ryckebusch *et al*, 2025; Westlake *et al*, 2024). The humoral arm is primarily mediated by the fat body and to a lesser extent by hemocytes, which respond to infection through the Toll and Imd pathways and produce antimicrobial peptides and other circulating immune effectors (Liegeois & Ferrandon, 2022). The cellular arm is mainly mediated by hemocytes, which perform phagocytosis, melanization-associated reactions, and encapsulation of larger invaders (Melcarne *et al*, 2019; Vlisidou & Wood, 2015). Among the different hemocyte populations, plasmatocytes represent the predominant phagocytic cell type that perform functions analogous to those of vertebrate macrophages (Gold & Brückner, 2014). These cells are therefore ideally positioned to provide mechanistic insight into the cellular basis of innate immune recognition and microbial clearance. In larvae, plasmatocytes are distributed across three main compartments: (i) the lymph glands, a reservoir that releases hemocytes upon parasitic infection; (ii) the circulating hemolymph; and (iii) sessile patches (Crozatier & Meister, 2007; Evans *et al*, 2003; Honti *et al*, 2010; Jung *et al*, 2005; Lanot *et al*, 2001; Makhijani *et al*, 2011; Makhijani & Brückner, 2012). Sessile hemocytes adhere to the inner surface of the larval body wall, forming clusters often closely associated with secretory cells known as oenocytes as well as with the termini of peripheral neurons. These cells dynamically exchange between sessile patches and the circulation (Babcock *et al*, 2008), and are mobilized from sessile sites into circulation upon parasitoid wasp infestation or mechanical stimulation of the cuticle, such as brushing (Márkus *et al*, 2009).

Investigations into phagocytosis have underscored the strong similarities between *Drosophila* and mammalian systems, highlighting the evolutionary conservation of this process (Melcarne *et al*, 2019; Ulvila *et al*, 2011). A large fraction of *Drosophila* phagocytic receptors belongs to the Nimrod family, which comprises 12 proteins characterized by specialized adhesive EGF-like domains known as NIM repeats (Somogyi *et al*, 2008). Nimrod family genes are predominantly clustered on the second chromosome and are classified into A-, B-, and C-type subgroups. A-type Nimrods (e.g., Draper and NimA) and C-type Nimrods (e.g., NimC1–4 and Eater) are transmembrane receptors involved in the engulfment of pathogens and apoptotic cells, in the maintenance of blood-brain barrier integrity as well as in regulating plasmatocyte adhesion and motility (Bretscher *et al*, 2015; Kocks *et al*, 2005; Kurant *et al*, 2008; Kurucz *et al*, 2007; Sakr *et al*, 2026). Among these, Eater and NimC1 are particularly important for bacterial phagocytosis (Melcarne *et al*, 2019; Kocks *et al*, 2005). In contrast, NimC4 (also referred to as SIMU) and Draper—a conserved member of the CED-1/MEGF10 family—recognize phosphatidylserine (PS) exposed on apoptotic cells (Freeman *et al*, 2003; Hamon *et al*, 2006; Kurant *et al*, 2008; MacDonald *et al*, 2006; Manaka *et al*, 2004; Mangahas & Zhou, 2005; Shklyar *et al*, 2013; Tung *et al*, 2013). Additional receptors, including integrins βν and αPS3, the CD36 receptor Santa-Maria (Hilu-Dadia *et al*, 2025; Nagaosa *et al*, 2011), and the bridging molecule Orion (Perron *et al*, 2023), have also been implicated in phagocytosis.

The five secreted B-type Nimrod proteins (NimB1–NimB5) encoded in the *Drosophila* genome remain less well characterized. Among them, NimB5 functions as a metabolic signal produced by the fat body, reducing hemocyte proliferation and adhesion during starvation (Ramond *et al*, 2020). By contrast, NimB4 and NimB1 exert opposing effects on efferocytosis, respectively enhancing or limiting this process upstream of Draper (Dolgikh *et al*, 2025; Logan *et al*, 2012; Manaka *et al*, 2004; Petrignani *et al*, 2021). Both proteins bind apoptotic cells via recognition of phosphatidylserine, acting as molecular bridging factors. Two NimB proteins encoded in the *Drosophila* genome, NimB2 and NimB3 remain uncharacterized. NimB2 is the most conserved member of the secreted NimB family across insects. In vitro studies have suggested that *Drosophila* NimB2 can bind very weakly to the Gram-negative bacterium *E. coli* (Zsámboki *et al*, 2013). Silencing of *NimB2* by injection of dsRNA has pointed to a role of this protein in the phagocytosis of *Staphylococcus aureus* but not of *Escherichia coli* bacteria in the mosquito *Anopheles gambiae* (Midega *et al*, 2013). As this function did not involve a direct interaction of AgNimB2 with *S. aureus*, it was proposed that AgNimB2 acts downstream of the thioester-containing protein (TEP) complement-like pathway activation, which promotes bacterial opsonization. The same study also reported an anti-*Plasmodium* effect of AgNimB2.

In this study, we characterized the function of *Drosophila* NimB2 and established its role as an opsonin that promotes the phagocytic clearance of *Staphylococcus aureus*. We show that NimB2 is produced by the fat body and secreted into the hemolymph, where it binds to *S. aureus* specifically and is required for effective control of *S. aureus* infection. Using bacterial cell wall mutants and binding assays, we provide evidence that NimB2 recognizes a lipoteichoic acid (LTA)–dependent structure on the *S. aureus* surface. Finally, we identify the phagocytic receptor Eater as a strong candidate hemocyte receptor for NimB2-coated bacteria. Together, these findings move beyond descriptive evidence of opsonin activity and define a molecular opsonin pathway in which fat-body-derived NimB2 coats *S. aureus* via an LTA-dependent interaction and bridges the bacterium to Eater on hemocytes, thereby promoting phagocytic clearance.

## Results

### NimB2 is expressed in the fat body and secreted into the hemolymph

To characterize the tissue-specific expression of *NimB2* in *Drosophila* larvae, we used a Trojan GAL4 fly line, that contains a cassette consisting of a *SA-T2A-GAL4-polyA* in the *NimB2* gene (referred to as *NimB2-Gal4*) allowing *Gal4* to recapitulate the endogenous expression pattern of *NimB2*. Consistent with FlyAtlas (Chintapalli *et al*, 2007), fluorescence of *NimB2-Gal4 > UAS-mCD8-GFP* larvae was detected predominantly in the fat body (Fig 1A and 1B), indicating that *NimB2* is mainly expressed in this tissue. This observation was supported by qPCR analysis of dissected larval tissues, which confirmed enriched *NimB2* transcript levels in the fat body of third-instar larvae (Appendix Fig S1A).

**Figure 1.**
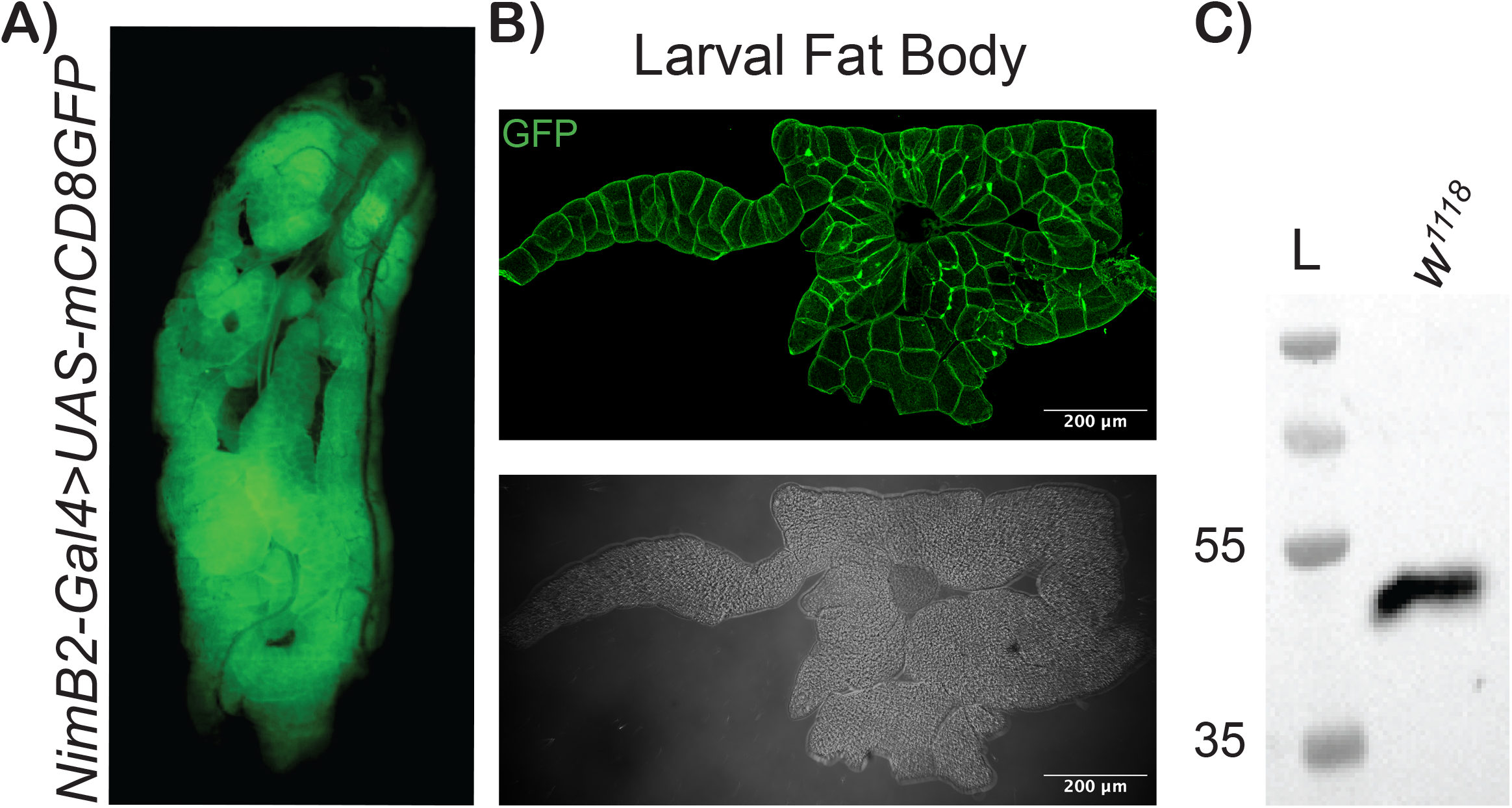
NimB2 is expressed in the fat body and secreted into the hemolymph. (A, B) Tissue-specific expression of *NimB2* in *Drosophila* larvae visualized using the *NimB2-Gal4 > UAS-mCD8-GFP* reporter system. (A) Representative image of a third-instar larva showing GFP expression. (B) Representative image of a dissected larval fat body showing strong GFP expression. (C) Western blot analysis of cell-free larval hemolymph using a rabbit anti-NimB2 antibody. The NimB2 protein was detected at approximately 50 kDa.

NimB2 carries a predicted N-terminal signal peptide and a recent proteomic analysis shows that NimB2 was the most abundant NimB protein in the hemolymph of flies (Rommelaere *et al*, 2025). To confirm that this protein is indeed secreted into the hemolymph, larval hemolymph was collected, hemocytes were removed by filtration, and the resulting cell-free fraction was analyzed by Western blot using a newly generated anti-NimB2 rabbit antibody. Detection of NimB2 from hemolymph samples (Fig 1C) confirmed that the protein is secreted, consistent with its predicted extracellular localization. Together, these observations establish NimB2 as a fat-body-derived secreted protein and support the hypothesis that it functions as a circulating immune factor.

### NimB2 mutant flies are specifically susceptible to S. aureus infection

To assess the role of NimB2 in antibacterial defense, we generated a viable *NimB2^Δ290^* mutant line by replacing a 292-bp fragment of the *NimB2* coding sequence with a 3xP3-DsRed cassette (Appendix Fig S1C) in the isogenic Drosdel background. Iso *NimB2^Δ290^* and wild-type (iso *w^1118^*) flies were challenged with a panel of Gram-negative and Gram-positive bacteria, including *Escherichia coli*, *Pectobacterium carotovorum carotovorum*, *Bacillus subtilis*, *Staphylococcus aureus* and *Enterococcus faecalis*. *Rel^E20^* flies, lacking a functional Imd pathway, were used as a susceptible control to Gram-negative bacteria while *Bom^Δ55c^* flies were used as a control for the Toll pathway response, as they are as susceptible as *spz^rm7^* flies when infected with Gram-positive bacteria (Lemaitre *et al*, 1996; Hedengren *et al*, 1999; Clemmons *et al*, 2015; Ryckebusch *et al*, 2025). *PPO1,PPO2 deficient* flies, that fail to melanize, were used as a positive control for *S. aureus* as they are known to rapidly succumb to this challenge (Binggeli et al., 2014). Among all the pathogens tested, *NimB2* mutants displayed a marked reduction in survival specifically after *S. aureus* infection (Fig 2A), whereas survival upon challenge with *E. coli*, *P. c. carotovorum*, *E. faecalis* and *B. subtilis* was comparable between mutants and wild-type (Appendix Fig. S2(A-D)), indicating that NimB2 selectively contributes to host defense against *S. aureus*. To reinforce our conclusion, we generated two independent null *NimB2* mutations, *NimB2^Δ49^*and *NimB2^Δ1122^*, each being caused by a CRISPR/Cas9-induced frameshift deletion (Appendix Fig S1D and E). Surprisingly, neither of the two mutant fly lines was viable while *NimB2^Δ49^/NimB2^Δ1122^* transheterozygous viable flies could be obtained. Importantly, *NimB2^Δ49^/NimB2^Δ1122^* flies also displayed a marked susceptibility to *S. aureus* as observed with the *NimB2^Δ290^* mutants (Appendix Fig S2E).

**Figure 2.**
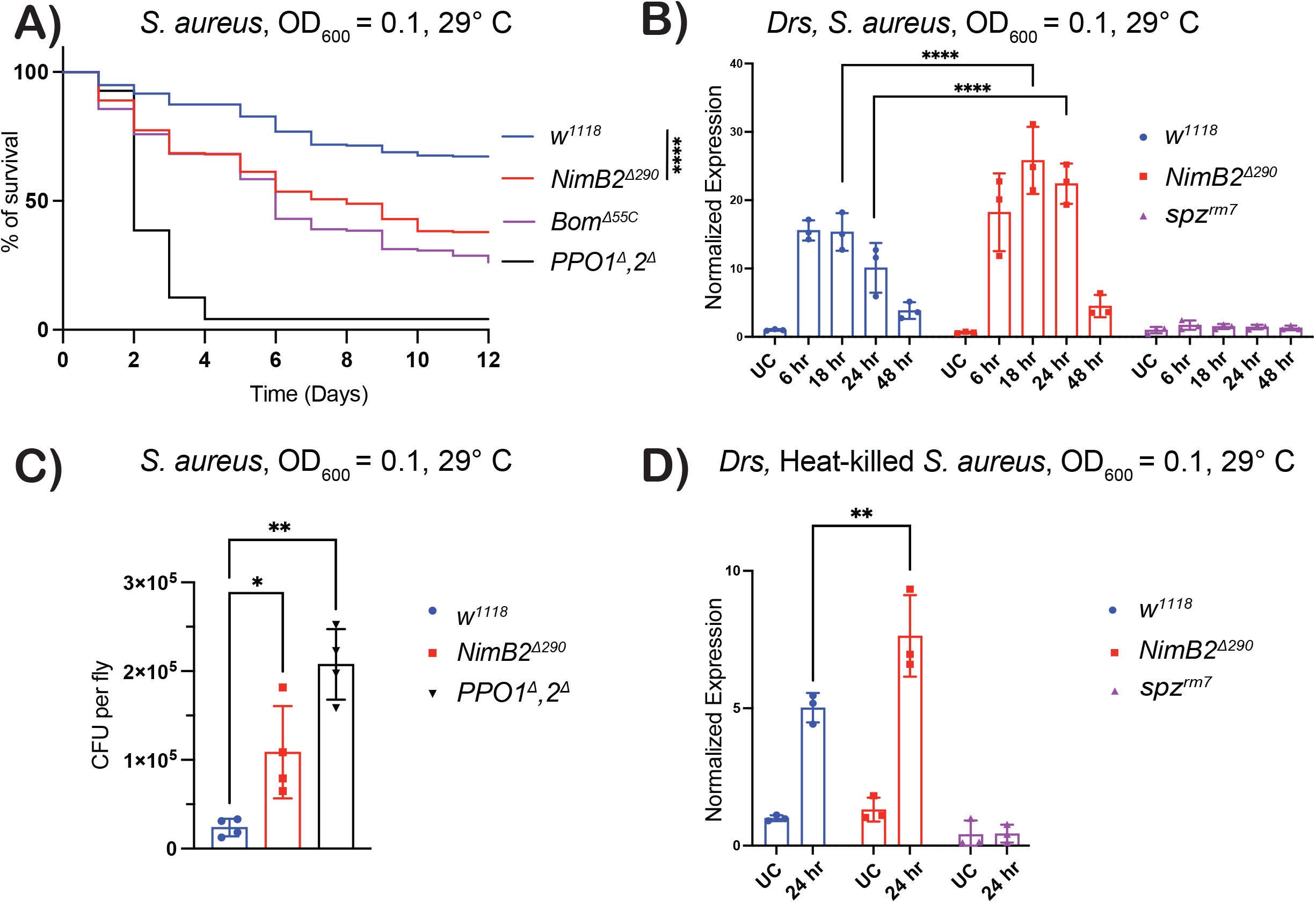
*NimB2* mutant flies are susceptible to *S. aureus* infection. (A) Survival of *wild-type* and *NimB2* mutant flies following systemic infection with *S. aureus*. Kaplan–Meier survival curve shows pooled data from three independent experiments, each consisting of three cohorts of 20 flies. Survival distributions were compared with *w^1118^* using the log-rank (Mantel–Cox) test. (B) *Drosomycin* expression following systemic infection with live *S. aureus*. (C) Bacterial load of wild-type and mutant flies measured by CFU assay 24 h after *S. aureus* infection. (D) *Drosomycin* expression following 24 h post-infection with heat-killed *S. aureus.* Each dot represents one biological repeat. Statistical significance was assessed using one-way analysis of variance (ANOVA). Statistical significance was defined as ns, not significant; P < 0.05 (*); P < 0.01 (**); P < 0.001 (***); and P < 0.0001 (****).

To determine whether this susceptibility reflected impaired humoral immunity, we monitored expression of the antifungal peptide gene *Drosomycin*, a readout of Toll pathway activation, following *S. aureus* infection. Upon systemic infection with *S. aureus*, *NimB2^Δ290^* mutants still induced *Drosomycin* expression, and transcript levels were even modestly elevated at 24 h post-infection compared with wild-type flies (Fig 2B), indicating that Toll-dependent antimicrobial peptide induction remains intact in *NimB2^Δ290^* mutant background. Because this increase could reflect a higher bacterial burden, we quantified bacterial load by colony-forming unit (CFU) assay 24 h after infection. *NimB2* mutant flies carried more *S. aureus* than wild-type controls (Fig 2C), supporting the idea that NimB2 limits bacterial proliferation and/or promotes bacterial clearance. We also measured *Drosomycin* expression following systemic injury with heat-killed *S. aureus* (Fig 2D). Under these conditions, *NimB2* mutants still displayed an higher *Drosomycin* expression compared to wild-type flies confirming that Toll pathway activation is intact in the absence of NimB2. The higher Toll pathway activation in *NimB2* mutants upon injection of dead *S. aureus* cannot be explained by increased bacterial growth, but could be due to a defect in the clearance of microbial elicitors that activate the pathway. Taken together, these results show that NimB2 is required for efficient control of *S. aureus* infection independently of Toll pathway activation.

### NimB2 is required for recognition and phagocytosis of *S. aureus* by hemocytes

Given the role of several Nimrod family members in phagocytosis, we asked whether NimB2 contributes to the uptake of *S. aureus* by hemocytes. *Ex vivo* phagocytosis assays using hemocytes from wandering third-instar larvae incubated with fluorescent *S. aureus* bioparticles revealed a reduction in phagocytic index in *NimB2^Δ290^* mutants compared with wild-type controls (Fig 3A). This phagocytic defect was less pronounced than that observed in hemocytes from *NimC1; eater* double-mutant larvae which are known to be deficient for phagocytosis of both Gram-negative and Gram-positive bacteria (Melcarne *et al*, 2019) (Fig 3A). Under the same conditions, loss of *NimB2* did not significantly affect phagocytosis of *E. coli* or apoptotic cell corpses, indicating that NimB2 preferentially promotes phagocytosis of *S. aureus* (Appendix Fig S3A and S3B). We further validated the *S. aureus* phagocytic defect phenotype with hemocytes from transheterozygous *NimB2^Δ49^/NimB2^Δ1122^* larvae (Appendix Fig S3C). To further assess the role of NimB2 in phagocytosis *in vivo*, we injected pHrodo-labeled *S. aureus* bioparticles into third-instar larvae, retrieved hemocytes 2 hours later and monitored the number of internalized particles. Hemocytes from *NimB2* mutants internalized fewer particles than hemocytes from wild-type larvae, while *NimC1; eater* double mutants again exhibited a pronounced uptake defect (Fig 3B and 3C). Similar results were obtained with a GFP-expressing *S. aureus* strain, further supporting a role for NimB2 in the efficient engulfment of *S. aureus in vivo* (Appendix Fig S3D and S3E). Together, these results indicate that NimB2 is required for optimal phagocytosis of *S. aureus* bioparticles.

**Figure 3.**
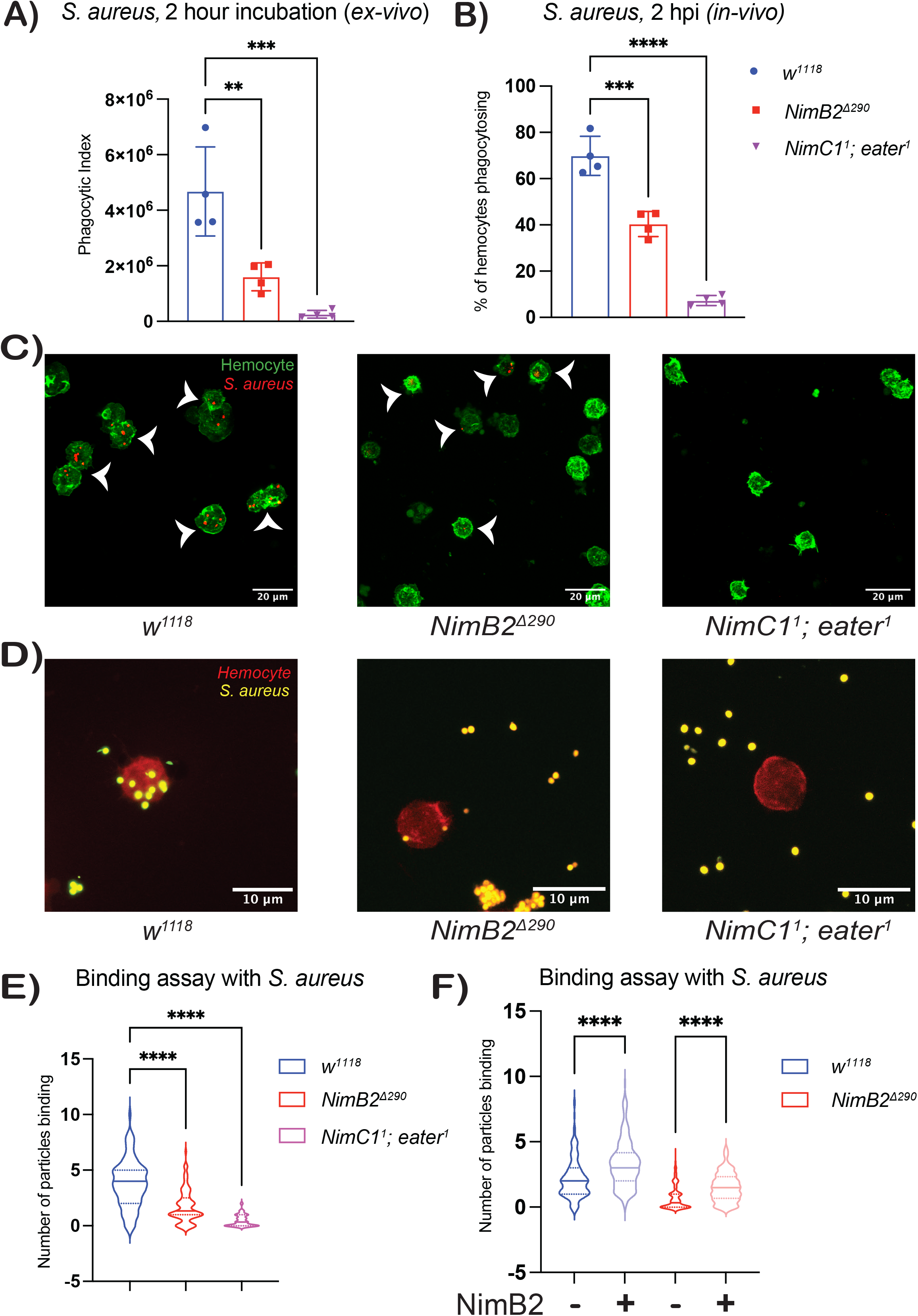
NimB2 is required for efficient recognition and phagocytosis of *S. aureus* by *Drosophila* hemocytes. (A) *Ex vivo* phagocytosis assay using larval hemocytes incubated for two hours with fluorescent *S. aureus* bioparticles. (B) *In vivo* phagocytosis assay monitoring the percentage of hemocytes phagocytosing pHrodo-labeled *S. aureus* bioparticles 2 hour after injection into third-instar larvae. Each dot represents one biological repeat. (C) Representative images from the *in vivo* phagocytosis assay showing hemocytes labeled with phalloidin (green) and pHrodo *S. aureus* bioparticles (red, arrow heads) with the indicated genotypes. (D) Representative image from cold-binding assay showing *S. aureus* bioparticles (yellow) attached to the surface of hemocytes labelled with phalloidin (red). *S. aureus* were incubated at 4°C for 1 hour with hemocyte from third-instar larvae. (E) Quantification of the cold-binding assay. (F) Addition of recombinant NimB2 protein (indicated by a ‘+’) enhances bioparticle binding in both *wild-type* and *NimB2* mutant backgrounds. Violin plot showing pooled data from three independent experiments. These data indicate that NimB2 acts at an early stage of phagocytosis by promoting bacterial recognition and binding. Statistical significance was assessed using one-way analysis of variance (ANOVA). Statistical significance was defined as ns, not significant; P < 0.05 (*); P < 0.01 (**); P < 0.001 (***); and P < 0.0001 (****).

The fact that NimB2 is a secreted protein suggested that it could act at an early stage of the phagocytic process. To define the step of phagocytosis controlled by NimB2, we examined the initial binding of *S. aureus* bioparticles to hemocytes using a cold-binding assay that allows particle attachment while minimizing internalization (Melcarne *et al*, 2019). Under these conditions, *NimB2* mutant hemocytes bound fewer *S. aureus* bioparticles than wild-type hemocytes (Fig 3D, and quantification in 3E). To test whether this defect could be rescued by extracellular NimB2, we produced a recombinant NimB2 protein fused to a human Fc–His tag (hFC-NimB2) in HEK cells. Because the Fc domain binds protein A on *S. aureus*, we removed the Fc-His tag by proteolytic cleavage to produce untagged NimB2 protein. In the following part of the manuscript, we referred to hFC-NimB2 for the tag version and NimB2 for the protein without the Fc tag. Addition of NimB2 protein increased the number of bound *S. aureus* bioparticles to both hemocytes from *wild-type* and *NimB2* third-instar larvae (Fig 3F). This rescue demonstrates that extracellular NimB2 is sufficient to enhance bacterial attachment to hemocytes indicating that NimB2 promotes the initial step of recognition and binding in the phagocytic defect. Taken together, the phagocytosis and binding assays support a model in which NimB2 functions as an opsonin that facilitates early recognition of *S. aureus* by hemocytes, thereby promoting efficient phagocytic clearance.

### Eater is required for NimB2-mediated recognition of *S. aureus*

Having established that NimB2 promotes the recognition and attachment of *S. aureus* to hemocytes, we next sought to identify the phagocytic receptor mediating this NimB2-dependent recognition. We examined several candidate phagocytic receptors previously implicated in the uptake of *S. aureus*, including Draper, Eater, Integrin-βν, and NimC1 (Kocks *et al*, 2005; Melcarne *et al*, 2019; Shiratsuchi *et al*, 2012). Hemocytes isolated from the corresponding mutant backgrounds were subjected to cold-binding assays using fluorescent *S. aureus* bioparticles in the presence or absence of recombinant NimB2. In wild-type hemocytes, addition of recombinant NimB2 markedly enhanced bacterial binding to hemocytes (Fig 4A). This enhancement was almost completely abolished in *eater* mutant hemocytes, whereas *draper*, *integrin-βν*, and *NimC1* mutant hemocytes displayed binding comparable to wild-type controls. These results identify Eater as the key receptor required for NimB2-dependent recognition of *S. aureus* by hemocytes and support a model in which NimB2 functions as an opsonin that promotes bacterial engagement by Eater.

**Figure 4.**
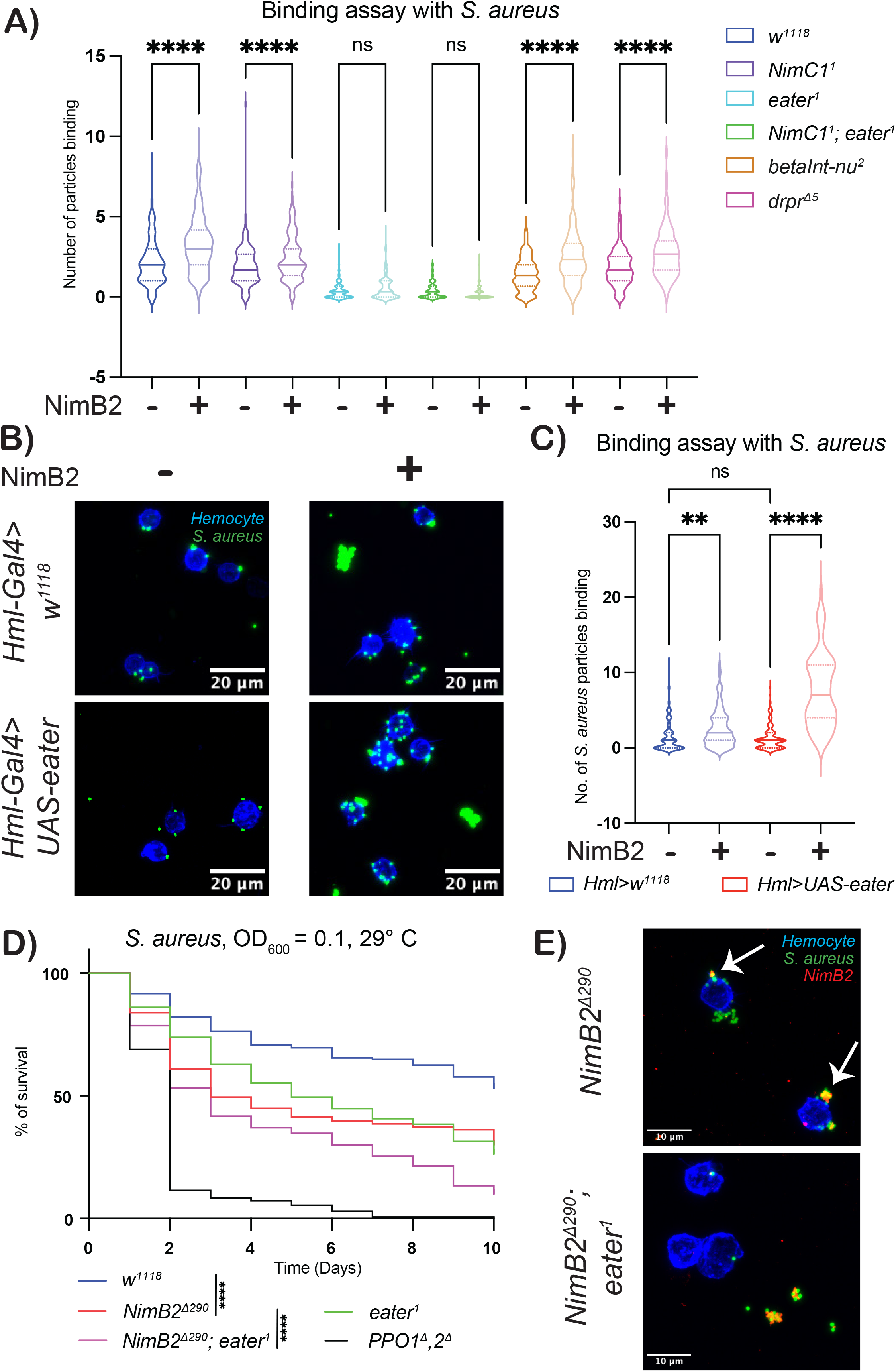
Eater is required for NimB2-mediated recognition of *S. aureus*. (A) Cold-binding assay using hemocytes from various phagocytic receptor mutant backgrounds in the presence (‘+’) or absence (‘-‘) of recombinant NimB2 and fluorescent *S. aureus* bioparticles. NimB2-dependent enhancement of bacterial binding was strongly reduced in *eater* mutant hemocytes but remained comparable to *wild-type* controls in the other receptor mutant backgrounds. (B) Representative confocal images from cold-binding assays with hemocytes from control (*Hml-Gal4>w^1118^*) and Eater-overexpressing (*Hml-Gal4>UAS-eater*) larvae incubated with *S. aureus* bioparticles in the absence (‘-‘) or presence (‘+’) of recombinant NimB2. Bioparticles (green) are associated with hemocytes labeled with phalloidin (blue). (C) Quantification of *S. aureus* bioparticle binding in control and Eater-overexpressing hemocytes in the presence or absence of recombinant NimB2. Violin plot showing pooled data from three independent experiments. Statistical significance was assessed using one-way analysis of variance (ANOVA). Statistical significance was defined as ns, not significant; P < 0.05 (*); P < 0.01 (**); P < 0.001 (***); and P < 0.0001 (****). (D) Survival of flies with the indicated genotypes following systemic infection with *S. aureus*. Kaplan–Meier survival curve shows pooled data from three independent experiments, each consisting of three cohorts of 20 flies. Survival distributions were compared with *w^1118^* using the log-rank (Mantel–Cox) test. The *NimB2; eater* double mutant showed significantly reduced survival compared with *NimB2* alone mutant (P<0.0001). (E) Representative image of binding assays using hemocytes from *NimB2* mutant and *NimB2*, *eater* double-mutant larvae incubated with *S. aureus* bioparticles preincubated with NimB2. In *NimB2* mutant, NimB2 (red) localizes to the bacterial surface (green) and is detected at the hemocyte–bacterium interface (arrow), consistent with a bridging role between *S. aureus* and hemocytes. Hemocytes were labeled with phalloidin (blue). In *NimB2, eater* double-mutant, NimB2 remains associated with the bacteria, but bioparticles fail to attach efficiently to the hemocyte surface, consistent with a requirement for Eater in productive recognition of NimB2-opsonized bacteria.

To further assess the contribution of *Eater*, we overexpressed *eater* in hemocytes using the *Hml-Gal4* driver and quantified bacterial binding in the presence of recombinant NimB2. Overexpression of *eater* significantly increased the association of *S. aureus* bioparticles with hemocytes compared with wild-type controls (Fig 4B and 4C), indicating that elevated Eater levels potentiate NimB2-mediated bacterial recognition. If the contribution of NimB2 to resist *S. aureus* infection were mediated exclusively through Eater, *NimB2; eater* double-mutant flies would be expected to phenocopy the reduced survival of *eater* single mutants following *S. aureus* infection. Instead, *NimB2; eater* double mutants showed a modest trend toward reduced survival following *S. aureus* infection compared with either single mutant (Fig 4D). This modest difference may reflect background effects or raises the possibility that NimB2 has additional antibacterial functions that are independent of Eater.

To further investigate the interaction between NimB2, *S. aureus*, and hemocytes, we performed confocal microscopy using hemocytes isolated from *NimB2* mutant and *NimB2; eater* double-mutant larvae. When *S. aureus* bioparticles were pre-incubated with recombinant NimB2 and added to *NimB2* mutant hemocytes, NimB2 localized to both the bacterial and the hemocyte membrane, consistent with a role as opsonin that favors attachment of bacteria to hemocytes. In contrast, although NimB2 remained associated with the bacterial, *S. aureus* bioparticles failed to bind efficiently to hemocytes from *NimB2; eater* double-mutant (Fig 4E). These observations indicate that NimB2 association with the bacterial surface is independent of Eater, whereas productive attachment of NimB2-opsonized bacteria to hemocytes requires Eater function. Collectively, these findings identify Eater as a key hemocyte receptor required for the recognition of NimB2-opsonized *S. aureus*.

### NimB2 selectively recognizes *S. aureus*

Having established that NimB2 promotes recognition of *S. aureus* by hemocytes, we next asked whether NimB2 binding is restricted to this species or extends to other bacteria. To address this, we then incubated recombinant NimB2 protein with different bacterial species. Following incubation, bacteria were collected by centrifugation, and the resulting pellets were resuspended in SDS sample buffer containing DTT and heated at 95°C. Samples were then separated by SDS-PAGE and analyzed by Western blot using a rabbit anti-NimB2 antibody to detect bacteria-associated NimB2. Under these conditions, NimB2 bound robustly to *S. aureus* but showed no detectable association with *Staphylococcus epidermidis*, *Staphylococcus saprophyticus*, *E. coli*, *E. faecalis*, *P. c. carotovorum* (*Ecc15*), or *B. subtilis* (Fig 5A). To exclude the possibility that the signal associated with *S. aureus* reflected Protein A–dependent detection, we confirmed that NimB2 also associates with *spa^-^ S. aureus* cells that are unable to produce Protein A. Of note, the *S. aureus* bioparticles (BioParticles^TM^, S23371) used in previous phagocytosis assays also lack Protein A, further indicating that NimB2 association with the bacterial surface is independent of Protein A. Microscopy-based analysis further confirmed the localization of NimB2 on the surface of *S. aureus*. For this assay, bacteria were incubated with recombinant NimB2, followed by detection using a rabbit anti-NimB2 primary antibody and an Alexa Fluor 488-conjugated secondary antibody. Fluorescence microscopy revealed a strong NimB2-associated signal on the surface of *S. aureus* cells (Fig 5C). Consistent with these observations, we further assessed NimB2 binding using an ELISA-based assay in which *spa^-^* bacterial cells were immobilized on microtiter plates and incubated with full-length recombinant hFC-NimB2. Bound NimB2 was detected using an HRP-conjugated goat anti-human IgG Fc antibody. This assay showed substantially higher NimB2 association with *S. aureus spa^-^* bacteria compared with the other bacterial species tested (Fig 5B). Together, these two approaches confirm that NimB2 directly associates specifically with the surface of *S. aureus* and that this interaction is selective among the bacterial species tested.

**Figure 5.**
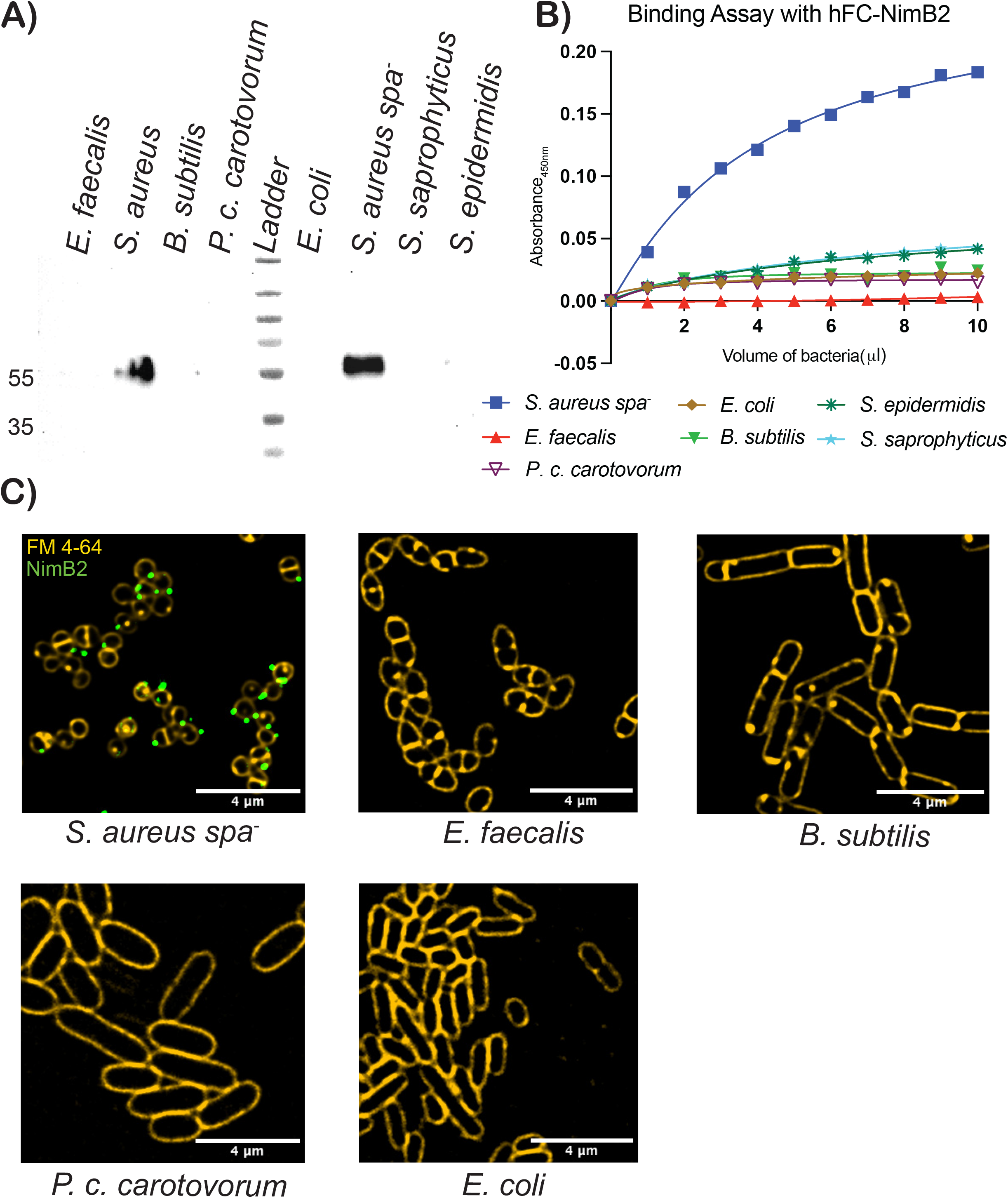
NimB2 selectively binds *Staphylococcus aureus*. (A) Western blot analysis of tag-cleaved recombinant NimB2 association with the indicated bacterial species using a rabbit anti-NimB2 antibody. (B) ELISA-based assay measuring the association of recombinant hFC-NimB2 with the indicated bacterial species. The binding curve shows pooled data from three independent experiments. (C) Representative fluorescence microscopy images showing bacterial association of NimB2. Bacterial cells were labeled with FM4-64 (yellow), and NimB2 was detected using a rabbit anti-NimB2 antibody (green).

### NimB2 bind to lipoteichoic acid associated determinant but not peptidoglycan

To identify the bacterial surface determinant recognized by NimB2, we first examined the contribution of two major *S. aureus* cell-surface components, peptidoglycan (PGN) and lipoteichoic acid (LTA). We initially tested the ability of recombinant hFC-NimB2 to bind commercially available preparations of *S. aureus* PGN and LTA by a dot-blot assay. As shown in Figure 6A, hFC-NimB2 bound strongly to the *S. aureus* PGN preparation from Sigma (77140), but showed only weak binding to the preparation from InvivoGen (tlrl-pgns2) and no detectable binding to the ultrapure InvivoGen preparation (tlrl-sipgn). Because commercial PGN preparations can contain residual cell-wall components, we assessed the presence of LTA in these PGN preparations using an anti-LTA antibody (Fig 6B). The Sigma PGN preparation, and to a lesser extent the InvivoGen preparation, contained detectable LTA, whereas no LTA was detected in the ultrapure InvivoGen PGN preparation. These results argue against PGN itself being the determinant recognized by NimB2. Strikingly, NimB2 binding closely correlated with the amount of contaminating LTA detected in the PGN preparations, suggesting LTA as a potential ligand.

**Figure 6.**
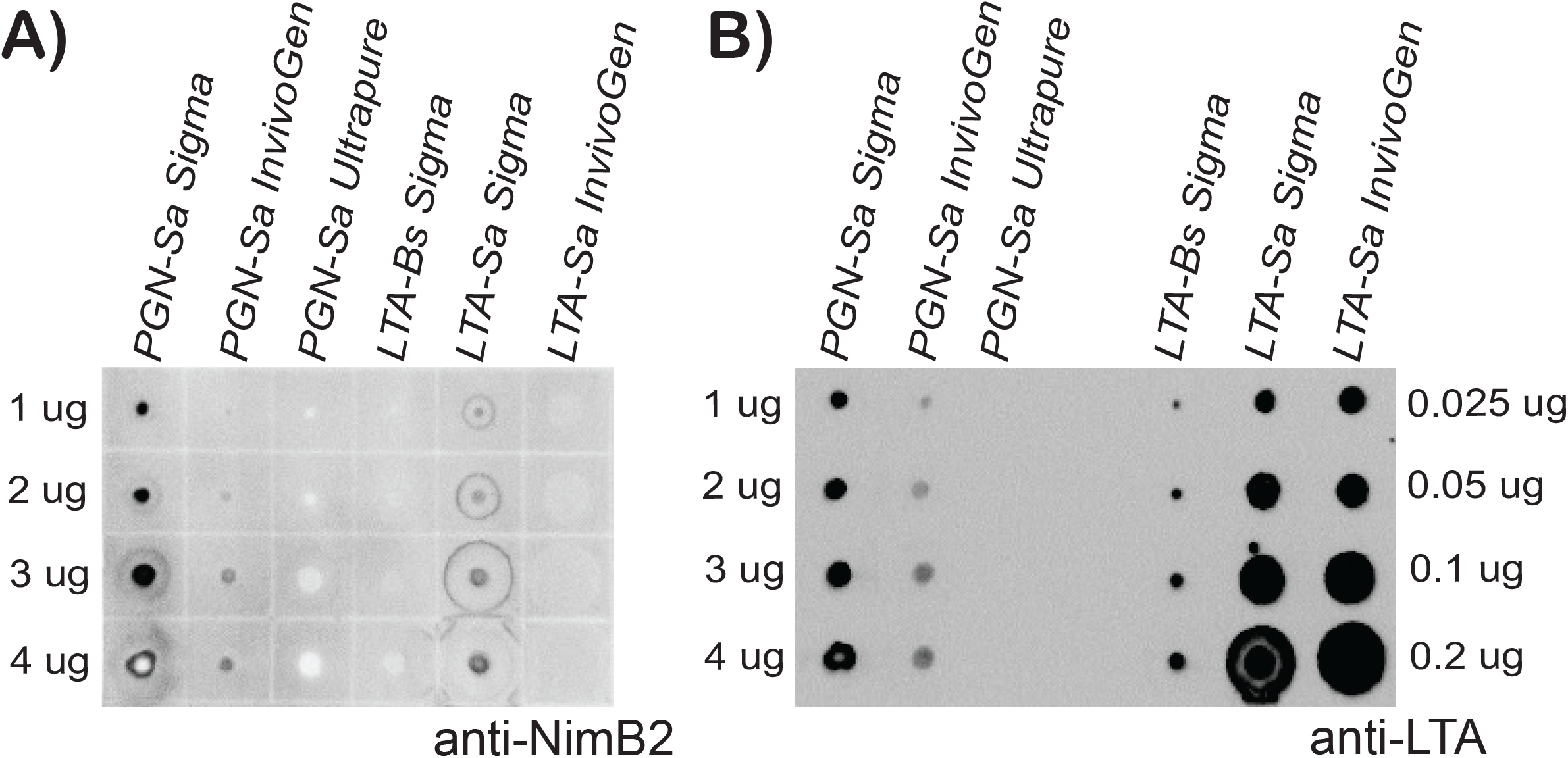
NimB2 preferentially associates with LTA-containing bacterial cell-wall preparations. (A) Dot-blot binding assay assessing NimB2 association with purified bacterial cell-wall components. Increasing amounts of peptidoglycan (PGN) from *S. aureus* (PGN-Sa Sigma, PGN-Sa InvivoGen, and PGN-Sa Ultrapure), Bacillus subtilis LTA (LTA-Bs Sigma), and S. aureus LTA (LTA-Sa Sigma and LTA-Sa InvivoGen) were spotted onto nitrocellulose membranes and incubated with recombinant NimB2. Bound NimB2 was detected using a rabbit anti-NimB2 antibody. The amounts of PGN and LTA spotted are indicated. (B) Dot-blot analysis of the same bacterial cell-wall preparations using an anti-LTA antibody to assess the presence and relative abundance of LTA in the preparations. The amounts spotted are indicated. NimB2 showed little or no detectable association with ultrapure *S. aureus* PGN, whereas stronger association was observed with LTA-containing preparations, supporting an LTA-dependent basis for NimB2 binding to the *S. aureus* cell surface.

Consistent with this hypothesis, we next directly tested LTA preparations from Sigma and InvivoGen. hFC-NimB2 bound to an *S. aureus* LTA preparation from Sigma (L2515), but not to an LTA preparation from *B. subtilis* (L3265). Interestingly, NimB2-hFc did not detectably bind to the *S. aureus* LTA preparation from InvivoGen (tlrl-pslta). The differential binding to the two *S. aureus* LTA preparations suggests that NimB2 does not recognize LTA solely on the basis of its presence, but may instead recognize a specific structural or modification state of LTA that can vary between preparations. Such differences could reflect variation in the composition, purification, or chemical modification of the LTA preparations. Together, the PGN and purified-component binding assays argue against PGN as the primary ligand and point to an LTA-dependent determinant on the *S. aureus* surface.

### Lipoteichoic acid and its D-alanylation modulate NimB2 binding to the *S. aureus* surface

To extent our analysis, we next examined the ability of NimB2 to bind to *S. aureus* mutants carrying mutations in enzymes involved in cell-wall biogenesis or remodeling. Because these mutants were generated in different *S. aureus* genetic backgrounds—spa-positive for USA300 and MW2, and spa-negative for NCTC—we used recombinant NimB2 without the Fc tag. Bacteria were incubated with NimB2, pelleted by centrifugation, and washed to remove unbound protein. The bacterial pellets were then treated with SDS to dissociate bound proteins, and the extracts were analysed by Western blot using an anti-NimB2 antibody. We first observed substantial differences in NimB2 binding among the three wild-type *S. aureus* strains, with markedly stronger binding to USA300 and MW2 than to NCTC (Fig 7A). We then compared NimB2 binding to two mutants in the NCTC background: (i) Δ*tagO* mutants, which are defective in wall teichoic acid (WTA) biosynthesis, and (ii) Δ*atl* mutants, which lack the major peptidoglycan hydrolase Atl and consequently accumulate increased amounts of peptidoglycan. As shown in Figure 7B, NimB2 bound more strongly to the *spa-* Δ*tagO* NCTC mutant than to the corresponding *spa^-^* wild-type strain. This result indicates that WTA is unlikely to be the determinant recognized by NimB2. Rather, the increased binding in the absence of WTA suggests that loss of this cell-wall component does not eliminate the NimB2-binding determinant but may increase NimB2 accessibility to another component, such as PGN or LTA.

**Figure 7.**
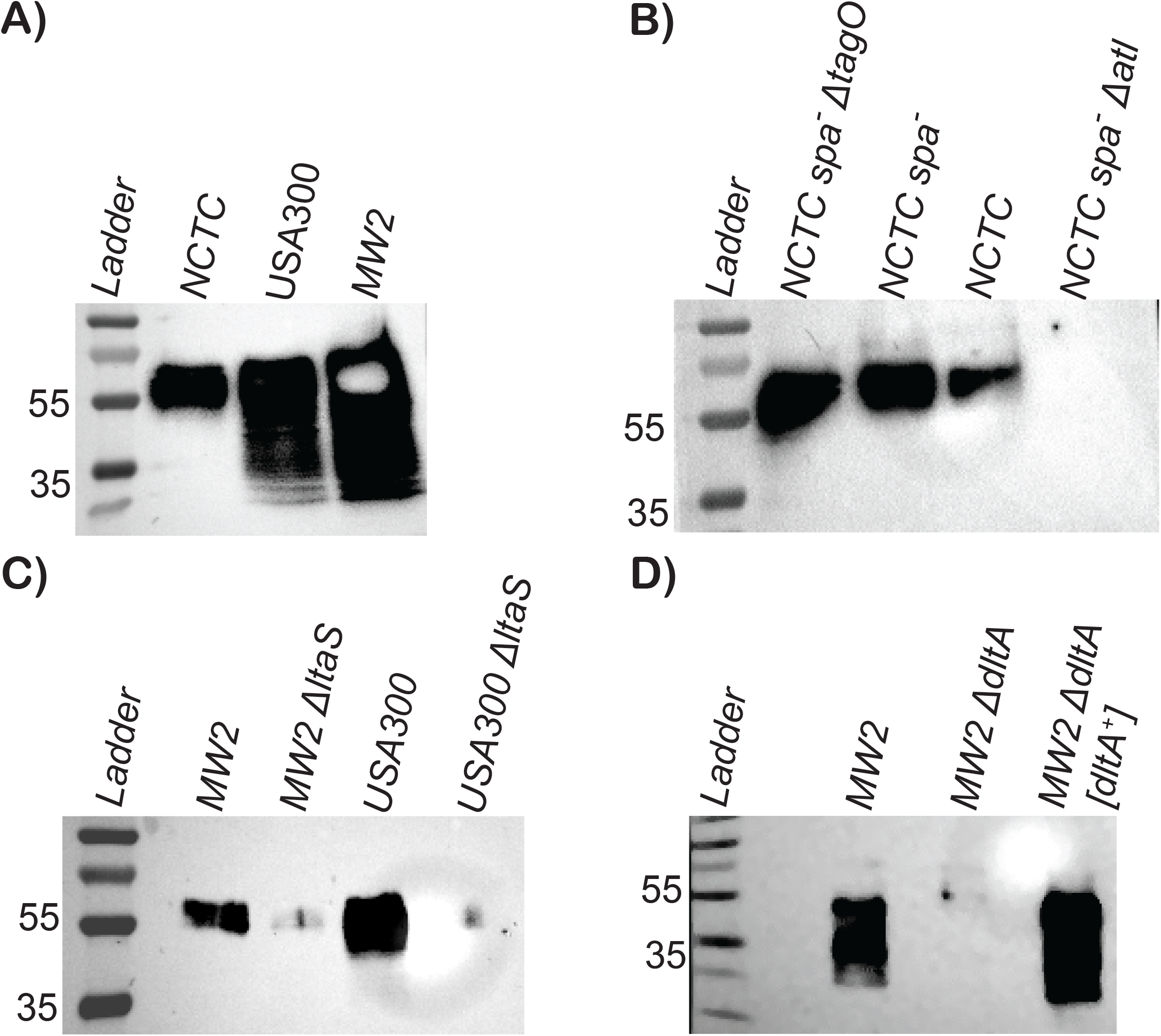
Lipoteichoic acid and its D-alanylation modulate NimB2 binding to the *S. aureus* surface. (A) Western blot analysis of NimB2 binding to *S. aureus* strains from the NCTC, USA300, and MW2 backgrounds, showing strain dependent differences in NimB2 association. (B) Western blot analysis of NimB2 binding to *spa−* NCTC mutants. NimB2 binding is increased in the *spa−* Δ*tagO* mutant and reduced in the *spa−* Δ*atl* mutant relative to the parental *spa−* strain. (C) Western blot analysis showing reduced NimB2 binding in Δ*ltaS* mutants in the MW*2* and USA300 backgrounds. (D) Western blot analysis of NimB2 binding to MW2 Δ*dltA* mutant and MW2 Δ*dltA* [dltA^+^] complemented strain.

Surprisingly, however, we detected no NimB2 binding to the Δ*atl* mutant, despite its increased exposure of PGN. Consistent with the Western blot analysis, microscopy-based analysis also showed reduced NimB2 association with the *Δatl* mutant (Appendix Fig S4A). Increased PGN exposure in Δ*atl* mutants has previously been demonstrated by the enhanced binding of the PGN-binding receptor PGRP-SA (Atilano *et al*, 2014). This result further supports our conclusion that PGN is unlikely to be directly recognized by NimB2. The absence of NimB2 binding to the Δ*atl* mutant could instead reflect reduced accessibility of NimB2 to an internal cell-wall component, such as LTA due to the excess of PGN. Together, these results further argue against PGN and WTA as the primary determinants recognized by NimB2 and are consistent with the possibility that NimB2 recognizes LTA or another cell-wall component whose accessibility is influenced by cell-wall architecture.

Finally, we analyzed the ability of NimB2 to bind to *S. aureus* Δ*ltaS* mutants lacking the lipoteichoic acid synthase. Strikingly, NimB2 binding was markedly reduced in Δ*ltaS* mutants, which are impaired in LTA synthesis, in two independent *S. aureus* strain backgrounds MW2 and USA300 (Fig 7C). Consistent with the Western blot analysis, an independent ELISA-based binding assay also showed reduced NimB2 association with the *ΔltaS* strains compared with their respective parental strains (Appendix Fig S4B). This result indicates that LTA is required for efficient association of NimB2 with the bacterial surface Interestingly, quantification of LTA levels in the three wild-type *S. aureus* strains revealed higher LTA abundance in USA300 and MW2 than in NCTC, paralleling the stronger NimB2 binding observed in these strains (Appendix Fig S4C). Collectively, these results indicate that NimB2 binding depends on LTA and suggest that LTA itself, or an LTA-associated determinant, may constitute the NimB2 ligand.

LTA can be modified notably the addition of D-alanine by Dlt enzymes, a modification that alter LTA charge and structure. Modification of LTA may explain the different ability of NimB2 to bind to various strains and LTA preparation. We therefore analyse the ability of NimB2 mutants to bind to Δ*dltA* mutants, which lack D-alanylation of teichoic acids. In the MW2 background, loss of *dltA* abolished NimB2 binding compared with the corresponding parental strain. We also observed that MW2 Δ*dltA* mutants complemented with a wild-type copy of *dltA* (*dltA [dltA^+^])* have wild-type NimB2 binding with MW2 (Fig 7D). These results were further supported by ELISA-based binding of NimB2 association with Δ*dltA* mutants (Appendix Fig S4D). These findings indicate that D-alanylation of LTA enhances NimB2 association with the bacterial surface, supporting a model in which NimB2 recognizes LTA itself or an LTA-dependent ligand whose affinity is modulated by *dltA*-dependent modifications.

## Discussion

Opsonins are soluble molecules that coat microbial or apoptotic surfaces and thereby greatly enhance their uptake by phagocytic cells (Flannagan *et al*, 2012; Rosales & Uribe-Querol, 2017; Wright & Douglas, 1904). By making targets more readily recognized by phagocytic receptors, they provide an additional layer of specificity and efficiency beyond direct receptor–ligand recognition. In vertebrates, antibodies and complement components are classical opsonins that promote pathogen clearance and strongly shape immune responses (Winkelstein, 1973). In insects, thioester-containing proteins (TEPs) have been proposed to play a similar role. Like complement factors, they carry an internal thioester motif, can covalently attach to microbial surfaces, and have been shown to promote phagocytosis in cell-based assays and *in vivo* infection models (Dostálová *et al*, 2017; Haller *et al*, 2018; Lagueux *et al*, 2000; Shokal & Eleftherianos, 2017). However, despite this complement-like activity, the molecular mechanisms by which TEPs and other insect opsonins recognize specific microbial ligands and connect to hemocyte receptors remain only partially understood. Studies in mosquitoes have shown that TEP1 is proteolytically processed into a two-chain form (TEP1-cut), exposing its reactive thioester and enables covalent deposition on pathogen surfaces. TEPs are often stabilized by other partners, such as LRIM1/APL1C (Fraiture *et al*, 2009; Povelones *et al*, 2009). In *Anopheles*, TEPs act through two main transmembrane receptors, LRP1 (LDL receptor-related protein 1) and BINT2 (β-integrin 2), each coupled to a distinct intracellular “engulfment” cascade (Moita *et al*, 2005). While there is strong evidence that *Drosophila* TEPs, notably TEP2, TEP3, and TEP4, contribute to bacterial phagocytosis (Dostálová *et al*, 2017; Haller *et al*, 2018; Stroschein-Stevenson *et al*, 2006), their mode of action and putative receptors remain poorly characterized. Secreted forms of Dscam were initially thought to promote phagocytosis, but recent data have challenged this claim (Westlake *et al*, 2026). Because some lectins and GNBPs are secreted proteins capable of binding microbial surfaces, they have been proposed as potential contributors to microbial opsonization; however, direct in vivo evidence that they function as opsonins to promote bacterial phagocytosis remains limited (Ao *et al*, 2007; Matskevich *et al*, 2010). Thus, while the fly immune system has been the focus of intense study, little is known about opsonins involved in bacterial phagocytosis.

In this study, we provide multiple lines of evidence supporting a role for NimB2 in the opsonization of bacteria. We show that NimB2 is expressed in the fat body, a major immune-responsive tissue, and secreted into the hemolymph. Analysis of *NimB2* null mutants reveals a role for NimB2 in host defense against *S. aureus*. While *NimB2* mutants display a wild-type humoral immune response, they have a reduced ability to phagocytose this bacterium. Using recombinant proteins, we show that NimB2 binds to *S. aureus* and promotes bacterial binding to hemocytes. Importantly, we demonstrate that NimB2 recognizes an Lipoteichoic acid (LTA)-dependent determinant on the *S. aureus* surface. It remains unclear whether NimB2 directly recognizes LTA itself or a factor conjugated to LTA. Several phagocytic receptors, notably Draper, Integrin-βν, and Eater, have been implicated in the uptake of *S. aureus*. Using loss-of-function mutations affecting each of the corresponding genes, we show that NimB2 mediates its effect specifically through Eater. Collectively, our results support a model in which fat body-derived NimB2 coats *S. aureus* by binding to an LTA-dependent surface determinant, thereby promoting bacterial internalization through the Eater receptor; thereby fulfilling the function of an opsonin (Fig 8B). A previous study also implicated NimB2 in the uptake of *S. aureus* in the mosquito *A. gambiae* (Midega *et al*, 2013), suggesting that the opsonizing function of NimB2 may be conserved across insects. However, that study did not identify either the receptor or the bacterial ligand involved. Moreover, AgNimB2 was reported not to bind *S. aureus* and was instead proposed to function downstream of activation of the Tep complement-like pathway (Midega et al., 2013). In contrast, our *in vitro* assays show that *Drosophila* NimB2 can bind *S. aureus* in the absence of other host factors. Future studies will be required to determine whether AgNimB2 can also bind *S. aureus* directly or whether it promotes phagocytosis through a distinct mechanism.

**Figure 8.**
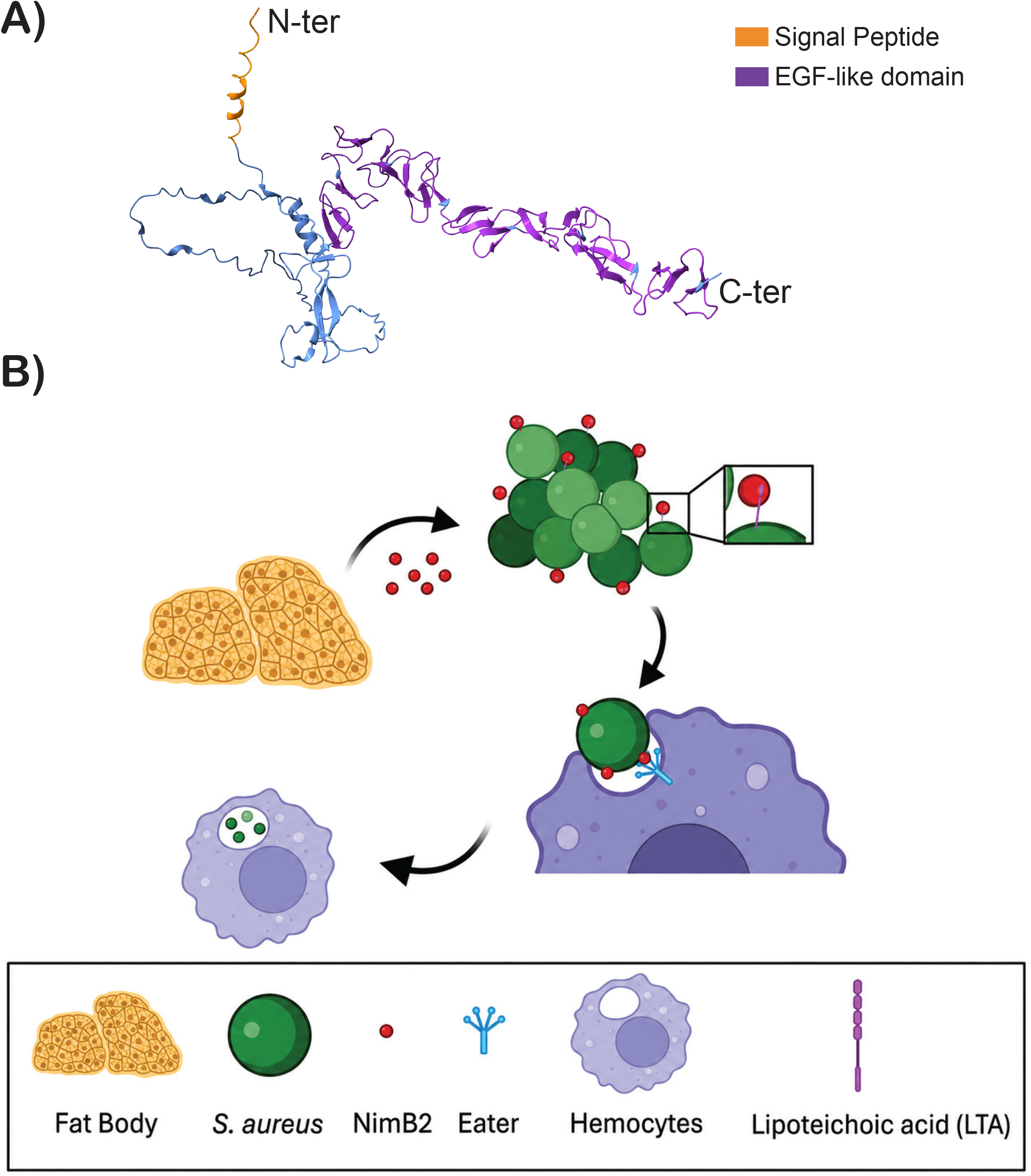
Proposed model for NimB2-mediated recognition of *S. aureus* by hemocytes. (A) Predicted domain organization of NimB2, showing the N-terminal signal peptide (orange) and EGF-like domain (purple) on the C-terminus. (B) Proposed model in which NimB2 is produced by the fat body and secreted into the hemolymph, where it binds to the surface of *S. aureus* through an LTA-dependent mechanism. D-alanylation of LTA, mediated by DltA, enhances NimB2 association with the bacterial surface. By coating the bacterial surface, NimB2 acts as a soluble opsonin that promotes bacterial recognition and attachment to hemocytes through the Eater receptor, thereby facilitating efficient phagocytic clearance.

On the hemocyte side, we identified Eater as a strong candidate receptor mediating NimB2-dependent recognition of *S. aureus*. Loss of *Eater* abolishes the NimB2-dependent enhancement of *S. aureus* binding to hemocytes, whereas *Eater* overexpression increases bacterial association in the presence of NimB2. Microscopy further indicates that NimB2 can bind bacteria in the absence of Eater, but that productive attachment to hemocytes requires Eater function. Although we have not yet demonstrated a direct biochemical interaction between NimB2 and Eater, the combined genetic and functional data provide strong support for a functional link between these two Nimrod proteins. An open question is whether NimB2 and Eater act exclusively in the same pathway or whether each protein has additional immune functions that are independent of the other. We favor the latter possibility. Eater has been shown to contribute to hemocyte sessility, whereas NimB2 has no detectable effect on the distribution of hemocytes in larvae (Appendix Fig S5B). In addition, Eater, together with NimC1, is involved in the uptake of Gram-negative bacteria (Melcarne *et al*, 2019). Whether NimB2 also contributes to the uptake of Gram-negative bacteria in cooperation with NimC1, remains to be investigated. Interestingly, we observed that *NimB2; eater* double mutants are slightly more susceptible to infection than either *NimB2* or *eater* single mutants. Although this increased susceptibility could result from a genetic background effect, we cannot exclude the possibility that NimB2 has functions beyond promoting phagocytosis. In support of this possibility, we found that NimB2 can promote the aggregation of *S. aureus* in the absence of Eater, a process that could potentially facilitate bacterial clearance through mechanisms independent of Eater-mediated phagocytosis.

The function of NimB2 appears to be very restricted to certain Gram-positive bacteria, such as *S. aureus*. However, *S. aureus* is not considered a natural pathogen of *Drosophila* (Broderick & Lemaitre, 2012; Cho & Kang, 2025). We hypothesize that NimB2 may contribute to host defense against other bacterial species encountered by flies in their natural environment that share surface features with *S. aureus*. Testing additional Gram-positive bacteria associated with flies may help clarify the physiological relevance and specificity of NimB2-mediated defense. It is noteworthy that the use of *S. aureus*, which is highly pathogenic to flies, has played a prominent role in *Drosophila* immunity research and has contributed to revealing the importance of several host defense mechanisms, including phagocytosis mediated by Eater, Draper, and β-integrin, as well as melanization (Binggeli *et al*, 2014; Hashimoto *et al*, 2009; Kocks *et al*, 2005; Shiratsuchi *et al*, 2012).

We investigated the molecular basis of NimB2 binding to the bacterial surface and found that NimB2 associates selectively with *Staphylococcus aureus*. Using *S. aureus* cell-wall mutants, we observed a marked reduction in NimB2 binding in Δ*ltaS* mutants, indicating that lipoteichoic acid (LTA) is required for efficient association of NimB2 with the bacterial surface. The finding that NimB2 binding to *S. aureus* depends on LTA is intriguing, as LTA is largely embedded within the cell envelope and lies beneath the wall teichoic acid and peptidoglycan layers of Gram-positive bacteria. AlphaFold modeling predicts that NimB2 (Jumper *et al*, 2021) consists of a signal peptide followed by two distinct regions: a short 65-amino-acid long, disordered N-terminal region and a larger, folded, β-sheet-rich domain with structural similarities to EGF, platelet, and complement domains (Fig 8A, Appendix S1F). The disordered region is enriched in polar residues, and is reminiscent of disordered regions found in cell-wall synthases associated with two-component systems that monitor cell-wall integrity in Gram-positive bacteria (Brogan *et al*, 2023, 2023, 2026). We speculate that this flexible N-terminal region could mediate binding to LTA, or potentially to another component of the cell envelope, as its flexibility may allow it to reach through pores in the peptidoglycan layer. In this model, the larger folded domain would remain exposed at the bacterial surface and could function as the opsonizing domain, facilitating recognition by Eater on hemocytes or promoting bacterial agglutination. At this stage, however, we cannot exclude the possibility that the binding to bacteria involved the EGF domain while the disordered domain contributes to receptor bridging. Interestingly, previous studies have suggested that Draper, best known for its role in efferocytosis (Manaka *et al*, 2004; Tung *et al*, 2013), can also bind LTA through its extracellular domain, and that this interaction contributes to the uptake of *S. aureus* (Hashimoto *et al*, 2009). Further studies will therefore be required to clarify the respective contributions of the NimB2/Eater and Draper pathways to the recognition and uptake of this bacterium. In mammals, LTA is also a potent inducer of innate immune responses and is recognized by several receptors, including TLR2/TLR6 heterodimers, with the contribution of secreted molecules such as CD14 and LBP (Kang *et al*, 2009; Schröder *et al*, 2003). Whether LTA recognition similarly contributes directly to the promotion of phagocytosis in mammals remains less well understood. Our findings therefore raise the possibility that LTA may serve not only as a microbial pattern recognized by innate immune receptors but also as a molecular cue for opsonization and phagocytic uptake.

The *dltABCD* operon encodes the enzymes responsible for adding D-alanine residues to both wall teichoic acid and lipoteichoic acid in the cell envelope of Gram-positive bacteria, including *S. aureus*, *Streptococcus pyogenes*, *Clostridioides* difficile as well as the *Drosophila* symbiont *Lactiplantibacillus plantarum* (Kristian *et al*, 2005; Matos *et al*, 2026; McBride & Sonenshein, 2011; Schneewind & Missiakas, 2014). D-alanylation reduces the net negative charge of the bacterial cell surface and thereby contributes to resistance to host innate immune effectors, including cationic antimicrobial peptides (Neuhaus & Baddiley, 2003). Loss of *dltA* markedly reduced NimB2 binding, indicating that D-alanylation of LTA enhances the interaction between NimB2 and the bacterial surface. This observation raises several interesting questions. Could differences in the extent or pattern of LTA D-alanylation contribute to the relatively restricted specificity of NimB2 for certain *Staphylococcus* species? How does D-alanylation promote NimB2 binding to the *S. aureus* surface? Addressing these questions may provide broader insights into how modifications of bacterial cell-wall components are sensed by the innate immune system and how such recognition can be coupled to phagocytic clearance.

In conclusion, our study identifies NimB2 as a secreted opsonin that promotes the phagocytic recognition of *S. aureus* in *Drosophila*. Together with the characterization of NimB1 and NimB4 as secreted factors acting upstream of Draper in efferocytosis (Dolgikh *et al*, 2025; Petrignani *et al*, 2021), our findings reveal an emerging role for secreted Nimrod proteins as bridging molecules/opsonins that connect targets to distinct phagocytic Nimrod receptors. This organization suggests that efficient phagocytosis in insects depends not only on the repertoire of receptors expressed by hemocytes, but also on secreted factors that determine how extracellular targets are recognized and presented to these receptors. The diversification of both secreted and phagocytic receptors of the Nimrod family may therefore have contributed to the broad and specialized phagocytic capabilities of the insect immune system. Since NimB2 is the most conserved secreted NimB, it is tempting to speculate that the first function of NimB was the opsonization of pathogens, and over evolutionary time, Nimod diversified in some insect lineage to other functions in efferocytosis such as NimB1 and NimB4 or metabolism such NimB5 (Dolgikh *et al*, 2025; Petrignani *et al*, 2021; Ramond *et al*, 2020). By defining both a bacterial determinant and a candidate phagocytic receptor for a secreted NimB protein, this study provides a molecular framework for understanding opsonin-mediated recognition in insects and opens new avenues for investigating the evolution and diversification of phagocytic immunity.

## Material and Methods

### *Drosophila* stocks, rearing, and mutant generation

All fly stocks used in this study and their corresponding references are listed in Table S1. Flies were maintained at 25°C on a 12 h light/dark cycle on standard fly medium containing 6% cornmeal, 6% yeast, 0.6% agar, and 0.1% fruit juice (50% grape juice and 50% multifruit/multivitamin juice), supplemented with 10.6 g L^-1^ Moldex and 4.9 mL L^-1^ propionic acid. Wandering third-instar larvae were collected at 110–120 h after egg laying (AEL).

The NimB2 null mutant *NimB2^Δ290^* was generated by replacing a 292-bp fragment in exon 2 with a 3xP3-DsRed cassette in an iso-*w^1118^* background, as shown in Appendix Fig S1C. Two additional frameshift alleles, *NimB2^Δ49^* and *NimB2^Δ1122^*, were generated by deleting 13 bp and 4 bp, respectively, in exon 1, as shown in Appendix Fig S1D and E. The fly lines carrying *NimB2^Δ49^* and *NimB2^Δ1122^* were lethal but *NimB2^Δ49^*/*NimB2^Δ1122^*flies were viable. The cause of this lethality is not known but could be an abberant NIimB2 toxic product or a nearby lethal mutation.

**Table S1:**
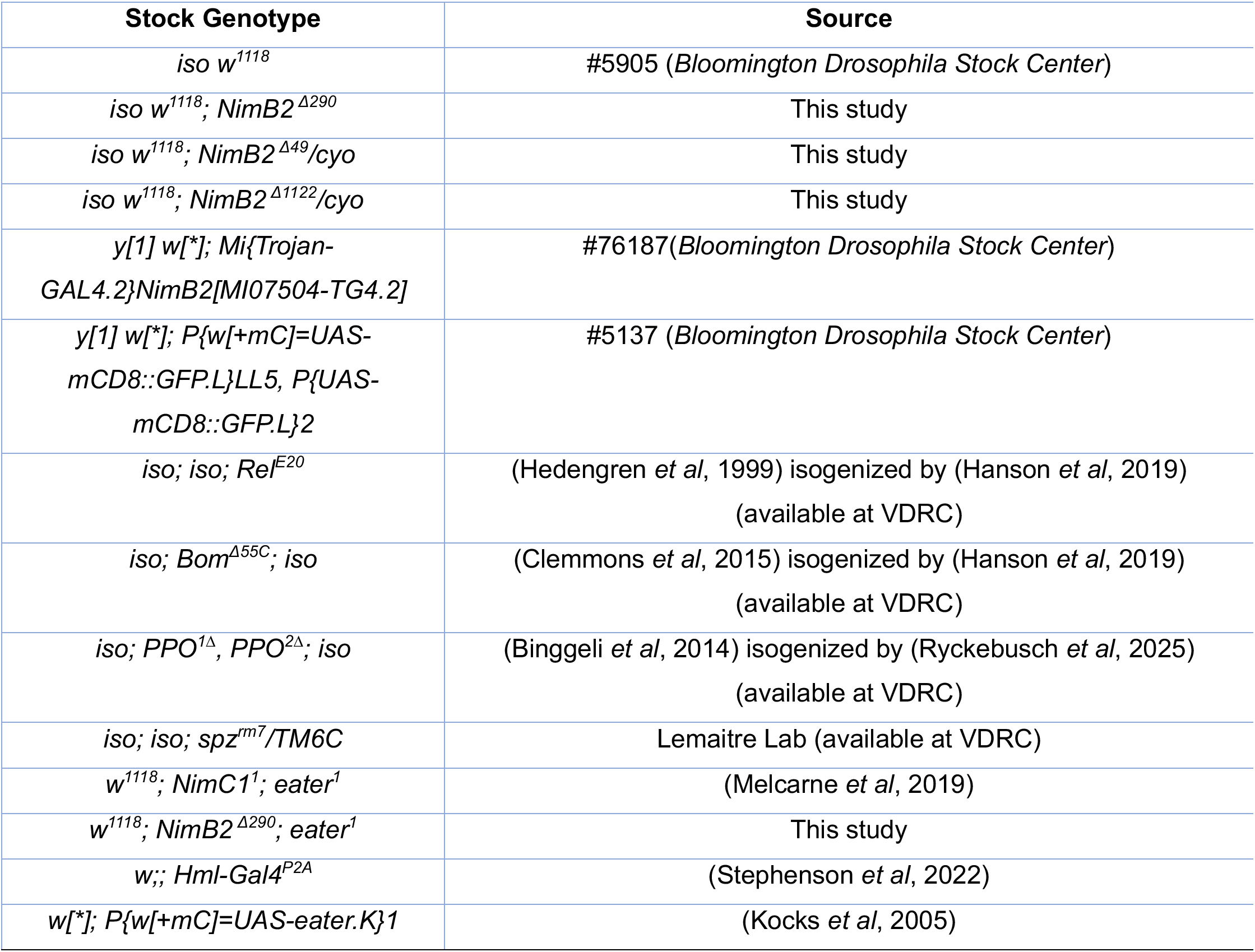
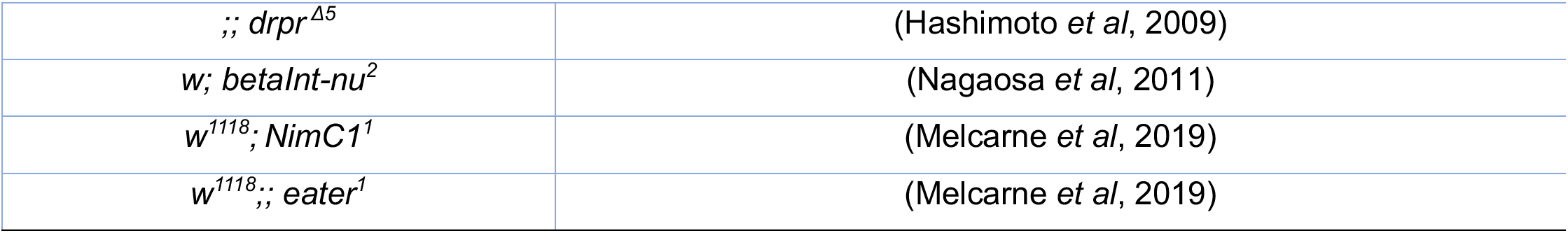
Genotypes of the fly lines used in this study.

### Bacterial stocks and culturing conditions

All bacterial strains used in this study and their corresponding references are listed in Table S2. *Staphylococcus aureus* strains, including the wild-type and mutant derivatives, as well as *Staphylococcus epidermidis*, *Staphylococcus saprophyticus*, *Bacillus subtilis*, *Escherichia coli*, and *Pectobacterium carotovorum carotovorum*, were cultured at 37°C in LB broth.

*Enterococcus faecalis* was cultured at 37°C in BHI medium.

Heat-killed microbes were prepared by two repeats of boiling at 95°C for 30 minutes then freezing at - 20°C for 30 minutes, before storage long term at -20°C. Microbe preparations were streaked onto agar plates to check for full efficiency of heat-killing before use in experiments.

**Table S2:**
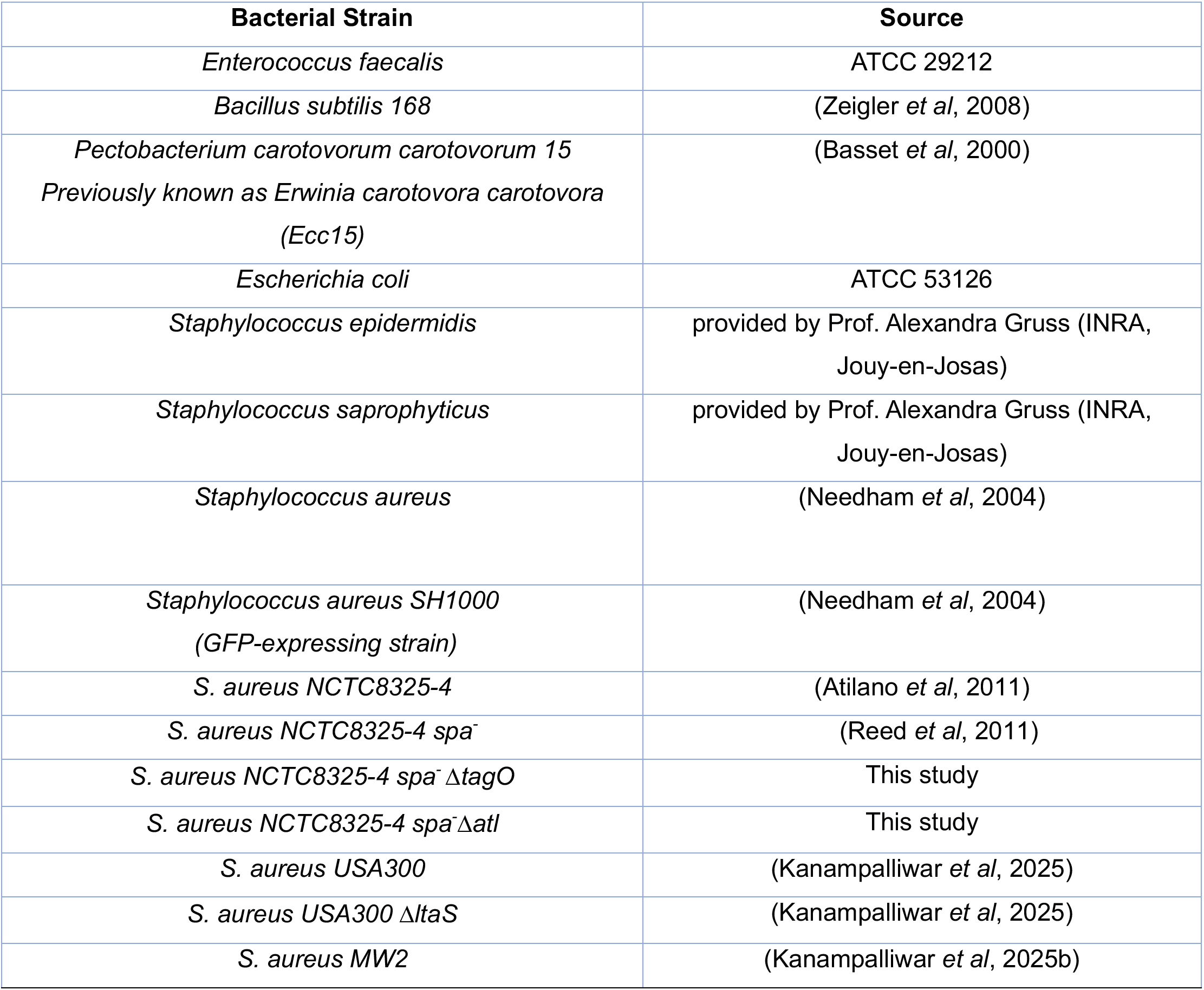
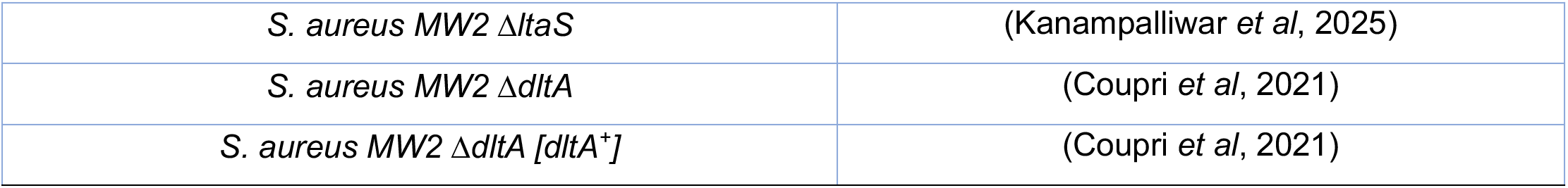
Bacterial strains used in this study.

### Recombinant NimB2 protein production and purification

Recombinant NimB2 protein was produced and purified with the assistance of the Protein Production and Structure Facility (PTPSP) at EPFL. NimB2 was expressed from the plasmid pcDNA3.1(+)uphos_ His-hFC-3C-NimB2, which encodes an N-terminal His-tagged human IgG Fc domain followed by a 3C protease cleavage site and NimB2. Recombinant protein was produced by transient transfection of 2 L HEK cells cultured in HyCell medium supplemented with valproic acid (VPA) to enhance recombinant protein expression. Six days after transfection, the culture supernatant was collected and incubated with MabSelect protein A resin (Cytiva/Amersham Biosciences, ref. 17-5199-03) for 4 h at 4°C with rotation. The resin was washed and bound protein was eluted using 0.1 M glycine, pH 3.0, followed by immediate neutralization with 1.5 M Tris, pH 8.0. The eluate was dialyzed overnight against PBS and subjected to 3C protease cleavage to remove the Fc-containing affinity tag. Following cleavage, the sample was incubated with MabSelect protein A resin (2 mL resin) for 2 h at 4°C to remove the cleaved Fc-containing fragment and any remaining uncleaved Fc-tagged NimB2 protein. The flow-through containing tag-cleaved NimB2 was concentrated using a 50-kDa molecular-weight-cutoff Amicon centrifugal filter. The concentrated NimB2 protein was further purified by size-exclusion chromatography on a Superdex 200 Increase 10/300 GL column equilibrated in PBS. Fractions containing NimB2 were pooled, and protein concentration was determined by absorbance at 280 nm. The purified protein was aliquoted in 1-mL volumes, flash-frozen, and stored at −80°C until use.

### Generation and purification of polyclonal anti-NimB2 antibody

Polyclonal antibodies against NimB2 were generated commercially (GenScript) using two New Zealand White rabbits. Recombinant untagged NimB2 protein was used as the immunogen. Prior to immunization, the antigen was validated by SDS-PAGE, with 5 mg of protein provided at a purity of >75% and a concentration of >0.4 mg/ml. Rabbits were immunized with recombinant untagged NimB2 using a three-injection Polyexpress immunization protocol (GenScript). The immunization phase was completed over approximately 4–6 weeks.

Following the third immunization, immune responses were evaluated by indirect ELISA. Both rabbits generated high anti-NimB2 immune titers. The antisera from the two rabbits were subsequently pooled and subjected to antigen-affinity chromatography for purification. A total of 40 ml of antiserum from each rabbit was used for pooled purification. The resulting antigen-affinity-purified polyclonal antibody was evaluated by indirect ELISA and stored as the final anti-NimB2 antibody.

### RT–qPCR experiments

For mRNA quantification, whole third-instar larvae (n = 15) or dissected tissues (n = 15) were collected and homogenized in TRIzol reagent according to the manufacturer’s protocol. Total RNA was extracted and resuspended in RNase-free MilliQ water. For infection experiments, flies were inoculated by pricking at the junction of the thoracic pleura with a needle dipped in a bacterial pellet and collected at 6, 18, 24, and 48 h post-infection. Samples were frozen at −20°C until RNA extraction. Total RNA was extracted from pooled samples of five flies using TRIzol reagent according to the manufacturer’s instructions. For reverse transcription, 500 ng of total RNA was converted to cDNA in a 10-µL reaction using PrimeScript RT (Takara) with a mixture of oligo(dT) and random hexamer primers. Quantitative PCR was performed using cDNA samples in 96-well plates on a LightCycler 480 (Roche) using SYBR Select Master Mix (Applied Biosystems) or PowerUp SYBR Green Master Mix (Applied Biosystems), as indicated for the respective experiments. Gene expression levels were normalized to RpL32.

Primers used in this study were:

Drs-F 5’- CGT GAG AAC CTT TTC CCA TAT GAT -3’

Drs-R 5’- TCC CAG GAC CAC CAG CAT -3’

NimB2-F 5’- TCA GTT CGT TGG CAA TGG CA -3’

NimB2-R 5’- GGC GGT AGA TGT GCG GAT TT -3’

RpL32-F: 5’-GAC GCT TCA AGG GAC AGT ATC TG-3’

RpL32-R: 5’-AAA CGC GGT TCT GCA TGA G-3’

### Systemic infections and survival assays

Systemic infections were performed by pricking 3- to 5-day-old adult males in the thorax with a 0.1 mm needle dipped into a concentrated bacterial pellet (Neyen *et al*, 2014; Troha & Buchon, 2019). At least two independent survival experiments were conducted for each infection. 20 infected flies were placed per vial and maintained on cornmeal fly medium containing 6.2 g agar, 58.8 g cornmeal, 58.8 g yeast, 60 mL grapefruit juice, 4.83 mL propionic acid, and 26.5 mL moldex (100 g L ^-1^ in ethanol) per liter with a flipping frequence of 2 days. Each replicate included three vials of 20 flies.

### Quantification of microbial load

3-5 days old flies were infected with the indicated microbe and concentration per OD_600_ as specified in the corresponding figure and allowed to recover. At the indicated time post-infection, flies were anesthetized using CO_2_ and surface sterilized by washing them in 70% ethanol. Ethanol was removed, and then flies were homogenized using a Precellys bead beater at 6500 rpm for 30 s in LB broth with 500 μL as pools of 5 flies. These homogenates were serially diluted and 100 µL was plated on LB agar plates. Plates were incubated overnight, and colony-forming units (CFUs) were counted using a Interscience Scan 500 plate scanning colony counter, and validated independently by manual counts for a subset of plates to ensure accuracy.

### Apoptotic cell preparation

S2 cells were cultured in Schneider’s insect medium (Sigma-Aldrich) supplemented with 10% fetal bovine serum (FBS; Gibco), penicillin (100 U/mL; Sigma-Aldrich), and streptomycin (100 U/mL; Sigma-Aldrich). Apoptosis was induced by adding cycloheximide (CHX; Sigma-Aldrich) to a final concentration of 50 µg/mL. After 24 h, cells were collected and pelleted by centrifugation at 400 × g for 5 min at 4°C. To label apoptotic bodies, the supernatant was incubated with CellTrace CFSE Cell Proliferation Kit (Invitrogen) or CellTracker Red CMTPX Dye (Invitrogen) at a final concentration of 5 µM for 15 min at room temperature in the dark.

### *Ex vivo* phagocytosis assay

Ex vivo phagocytosis assays were performed as described by (Melcarne *et al*, 2019). Briefly, five third-instar larvae were surface sterilized and bled into 120 µL Schneider’s insect medium within 2 min. Hemolymph was collected in Protein LoBind Eppendorf tubes to minimize sample loss. Approximately, 2 × 10^4^ apoptotic bodies or 10^5^ bacterial bioparticles (E13231, S23372, Invitrogen) were added to the hemocyte suspension and incubated for 2 h at room temperature. Samples were then placed on ice and immediately analyzed by flow cytometry using a CytoFLEX system (Beckman Coulter).

The phagocytic index was calculated according to the following equations:

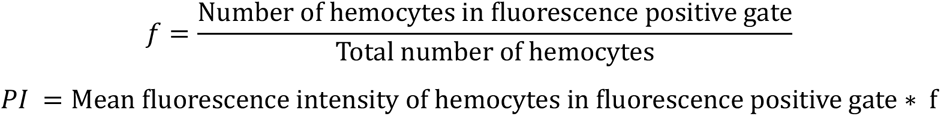

### *In vivo* phagocytosis assay

For *in vivo* phagocytosis assays, approximately 10^4^ bacterial bioparticles (A10010) or live bacterial cells were injected into wandering third-instar larvae using a Nanoject microinjector. Injections were performed laterally into the posterior hemocoel to promote systemic distribution. Larvae were then maintained at 25°C for 2 h to allow phagocytosis to proceed under physiological conditions. After incubation, larvae were washed in 1× PBS, dissected in Schneider’s insect medium, and hemocytes were collected onto glass slides.

Hemocytes were fixed in 4% paraformaldehyde (PFA) for 15 min at room temperature, washed three times with 1× PBS to remove unbound bacteria and debris, and imaged by confocal microscopy using an Olympus FV4000 system. For quantification, at least 100 hemocytes were scored per condition in each experiment, and the percentage of hemocytes containing fluorescent bacterial particles was determined using Fiji.

### Cold-binding assay

To assess bacterial binding, hemocytes were isolated by bleeding wandering third-instar larvae into 120 µL cold Schneider’s insect medium on pre-chilled glass slides. Approximately 10^5^ live fluorescent bacteria were added immediately after bleeding, and samples were incubated on ice for 60 min to permit binding while minimizing internalization. For rescue experiments, recombinant NimB2 protein was added together with the bacterial bioparticles at a final concentration of 10 µM.

Hemocytes were fixed in 4% PFA for 15 min at room temperature. After fixation, cells were washed three times with 1× PBS to remove unbound bacteria and debris and imaged using an Olympus FV4000 confocal microscope with a 60× objective. Image analysis was performed in Fiji, and the number of bound bacteria per hemocyte was quantified by scoring at least 100 hemocytes per genotype in each experiment.

### Protein-bacteria binding assay

For bacterial binding assays, bacterial cultures were first adjusted to an OD_600_ of 1.0 and collected by centrifugation. The bacterial pellets were washed and resuspended in PBS to remove the culture medium. Tag-cleaved recombinant NimB2 protein was then added to the bacterial suspension at a final concentration of 100 nM. Samples were incubated at 25°C for 1 h to allow NimB2 to associate with the bacterial surface. Bacteria were subsequently collected by centrifugation at 8000 rpm for 15 min and washed three times with PBS to remove unbound protein. The resulting bacterial pellets were analyzed by Western blotting or fluorescence microscopy to assess NimB2 association with the bacterial surface.

#### Western blot–based binding assay

For Western blot analysis, bacterial pellets obtained following the binding assay were resuspended in 4× SDS sample buffer, boiled at 95°C for 5 min, and separated by SDS-PAGE on a 4%–12% precast Novex NuPAGE gel (Invitrogen) under reducing conditions, and proteins were transferred to membranes using an iBlot 2 system (Invitrogen). Membranes were blocked in 5% non-fat dry milk in PBS containing 0.1% Tween 20 for 1 h, then incubated overnight at 4°C with rabbit anti-NimB2 antibody (GenScript; 1:1,000). After washing, membranes were incubated with HRP-conjugated anti-rabbit secondary antibody (Jackson ImmunoResearch; 1:20,000) for 45 min at room temperature. Bound antibody was detected using enhanced chemiluminescence (ECL; GE Healthcare) according to the manufacturer’s instructions, and membranes were imaged using a ChemiDoc XRS+ system (Bio-Rad).

#### Microscopy-based binding assay on bacteria

For microscopy-based binding assays, bacteria were incubated with rabbit anti-NimB2 primary antibody (1:1,000) for 1 h at 25°C, followed by incubation with Alexa Fluor 488-conjugated anti-rabbit secondary antibody (1:2,000). Bacterial membranes were counterstained with FM 4-64 (1:200) for 45 min at 25°C. Samples were mounted on agar pads, covered with coverslips, and imaged using a Nikon-CrestOptics SIM microscope equipped with a 100× objective. Images were analyzed with Fiji.

### ELISA-based binding assay

For ELISA-based binding assays, serial volumes of bacterial suspensions adjusted to OD_600_ of 1 were added to binding plates and incubated at room temperature for 1.5 h. Bacteria were fixed with 4% paraformaldehyde for 15 min, and wells were blocked overnight at 4°C with blocking buffer. Wells were subsequently incubated with full-length recombinant NimB2 (hFC-NimB2) for 30 min at room temperature at a final concentration of approximately 100 nM per well. After washing, bound NimB2 was detected using HRP-conjugated goat anti-human IgG Fc antibody (1:15,000) for 1 h. Color development was performed using 1-Step Slow TMB-ELISA substrate (50 µL per well) for 30 min or until color developed, and the reaction was stopped by adding 2 M sulfuric acid (50 µL per well) for 5 min. Absorbance was measured at 450 nm using a plate reader.

### Dot-blot binding assay

For the NimB2-binding assay, nitrocellulose membranes were spotted with 1, 2, 3, and 4 µg of the indicated samples (both PGN and LTA). The membranes were allowed to dry for 30 min at room temperature. Membranes were then blocked overnight at 4°C in 5% BSA and subsequently incubated in SuperBlock buffer (Thermofisher^TM^ 37536) for 10 min at room temperature. Membranes were then incubated with full-length recombinant hFc-NimB2 protein (75 µL of a 1 mg/mL hFc-NimB2 protein stock diluted in 20 mL SuperBlock buffer, corresponding to a final protein concentration of 3.75 µg/mL) for 30 min at room temperature. Membranes were washed four times for 5 min each with PBS Tween (PBST) 0.1% and incubated with HRP-conjugated goat anti-human IgG Fc antibody (1:15,000) for 1 h at room temperature. Following four additional 5-min washes with PBST, signal was detected using ECL substrate and imaged using a Bio-Rad^TM^ imaging system.

For detection of LTA, nitrocellulose membranes spotted with PGN or LTA were blocked overnight at 4°C in 5% BSA and subsequently incubated in SuperBlock buffer for 10 min. at room temperature. Membranes were incubated with mouse anti-LTA antibody (MA1-7402) (1:400; 50 µL antibody in 20 mL SuperBlock buffer) for 1 h at room temperature. After four washes of 5 min each with PBST, membranes were incubated with HRP-conjugated anti-mouse secondary antibody (1:10,000; 4 µL in 40 mL SuperBlock buffer) for 1 h at room temperature. Membranes were washed four additional times with PBST, and signal was detected using ECL substrate and imaged using a Bio-Rad^TM^ imaging system.

### Statistical analysis

All experiments were performed with at least three independent biological replicates unless stated otherwise. Statistical analyses were conducted using GraphPad Prism version 9. Unless otherwise specified, each point in the graphs represents one biological replicate. Statistical significance was assessed using an unpaired Student’s t test for pairwise comparisons and one-way analysis of variance (ANOVA) for comparisons involving more than two groups. For survival analyses, Kaplan–Meier survival curves were generated and survival distributions were compared using the two-sided log-rank (Mantel–Cox) test. Statistical significance was defined as ns, not significant; P < 0.05 (*); P < 0.01 (**); P < 0.001 (***); and P < 0.0001 (****).

## Acknowledgments

We thank Dominik Alwardt for teaching us the microscopy techniques used to study bacteria– protein interactions and Prof. Mariana Gomes de Pinho for hosting Prince in her research laboratory. We are grateful to the BIOP platform (EPFL) for their assistance with microscopy, the PTSPF platform (EPFL) for producing the recombinant NimB2 protein, and the PTCF platform (EPFL) for their support with FACS experiments. We also thank Prof. Taeok Bae and Prof. Aurélie Budin-Verneuil for kindly providing bacterial strains. We thank Prof. Andrea David Vettiger and Prof. Josué Flores Kim for their valuable expertise and advice on bacterial biology. We are grateful to the Bloomington *Drosophila* Stock Center (BDSC) and Vienna *Drosophila* Stock Center (VDRC) for providing some of the *Drosophila* stocks used in this study.

## References

Ao J, Ling E & Yu X-Q (2007) Drosophila C-type Lectins Enhance Cellular Encapsulation. Mol Immunol 44: 2541–2548

Atilano ML, Pereira PM, Vaz F, Catalão MJ, Reed P, Grilo IR, Sobral RG, Ligoxygakis P, Pinho MG & Filipe SR (2014) Bacterial autolysins trim cell surface peptidoglycan to prevent detection by the Drosophila innate immune system. eLife 3: e02277

Atilano ML, Yates J, Glittenberg M, Filipe SR & Ligoxygakis P (2011) Wall Teichoic Acids of Staphylococcus aureus Limit Recognition by the Drosophila Peptidoglycan Recognition Protein-SA to Promote Pathogenicity. PLOS Pathog 7: e1002421

Babcock DT, Brock AR, Fish GS, Wang Y, Perrin L, Krasnow MA & Galko MJ (2008) Circulating blood cells function as a surveillance system for damaged tissue in Drosophila larvae. Proc Natl Acad Sci U S A 105: 10017–10022

Basset A, Khush RS, Braun A, Gardan L, Boccard F, Hoffmann JA & Lemaitre B (2000) The phytopathogenic bacteria Erwinia carotovora infects Drosophila and activates an immune response. Proc Natl Acad Sci U S A 97: 3376–3381

Binggeli O, Neyen C, Poidevin M & Lemaitre B (2014) Prophenoloxidase Activation Is Required for Survival to Microbial Infections in Drosophila. PLOS Pathog 10: e1004067

Bloomington Drosophila Stock Center Bloomingt Drosoph Stock Cent

Bretscher AJ, Honti V, Binggeli O, Burri O, Poidevin M, Kurucz É, Zsámboki J, Andó I & Lemaitre B (2015) The Nimrod transmembrane receptor Eater is required for hemocyte attachment to the sessile compartment in Drosophila melanogaster. Biol Open 4: 355–363

Broderick N & Lemaitre B (2012) Gut-associated microbes of Drosophila melanogaster. Gut Microbes 3: 307–321

Brogan AP, Habib C, Hobbs SJ, Kranzusch PJ & Rudner DZ (2023) Bacterial SEAL domains undergo autoproteolysis and function in regulated intramembrane proteolysis. Proc Natl Acad Sci 120: e2310862120

Brogan AP, Schmid EW & Rudner DZ (2026) A broadly conserved gram-positive lipoprotein regulates cell elongation. Proc Natl Acad Sci 123: e2610431123

Chintapalli VR, Wang J & Dow JAT (2007) Using FlyAtlas to identify better Drosophila melanogaster models of human disease. Nat Genet 39: 715–720

Cho KH & Kang SO (2025) The Gut Microbiota of Drosophila melanogaster: A Model for Host-Microbe Interactions in Metabolism, Immunity, Behavior, and Disease. Microorganisms 13: 2515

Clemmons AW, Lindsay SA & Wasserman SA (2015) An effector Peptide family required for Drosophila toll-mediated immunity. PLoS Pathog 11: e1004876

Coupri D, Verneuil N, Hartke A, Liebaut A, Lequeux T, Pfund E & Budin-Verneuil A (2021) Inhibition of d-alanylation of teichoic acids overcomes resistance of methicillin-resistant Staphylococcus aureus. J Antimicrob Chemother 76: 2778–2786

Crozatier M & Meister M (2007) Drosophila haematopoiesis. Cell Microbiol 9: 1117–1126

Diao F, Ironfield H, Luan H, Diao F, Shropshire WC, Ewer J, Marr E, Potter CJ, Landgraf M & White BH (2015) Plug-and-Play Genetic Access to *Drosophila* Cell Types using Exchangeable Exon Cassettes. Cell Rep 10: 1410–1421

Dolgikh A, Rommelaere S, Ghanem A, Petrignani B, Poidevin M, Kurant E & Lemaitre B (2025) The secreted Nimrod protein NimB1 negatively regulates early steps of apoptotic cell phagocytosis in Drosophila. Development 152: dev204919

Dostálová A, Rommelaere S, Poidevin M & Lemaitre B (2017) Thioester-containing proteins regulate the Toll pathway and play a role in Drosophila defence against microbial pathogens and parasitoid wasps. BMC Biol 15: 79

Evans CJ, Hartenstein V & Banerjee U (2003) Thicker than blood: conserved mechanisms in Drosophila and vertebrate hematopoiesis. Dev Cell 5: 673–690

Flannagan RS, Jaumouillé V & Grinstein S (2012) The cell biology of phagocytosis. Annu Rev Pathol 7: 61–98

Fraiture M, Baxter RHG, Steinert S, Chelliah Y, Frolet C, Quispe-Tintaya W, Hoffmann JA, Blandin SA & Levashina EA (2009) Two Mosquito LRR Proteins Function as Complement Control Factors in the TEP1-Mediated Killing of *Plasmodium*. Cell Host Microbe 5: 273–284

Freeman MR, Delrow J, Kim J, Johnson E & Doe CQ (2003) Unwrapping glial biology: Gcm target genes regulating glial development, diversification, and function. Neuron 38: 567–580

Gold KS & Brückner K (2014) *Drosophila* as a model for the two myeloid blood cell systems in vertebrates. Exp Hematol 42: 717–727

Haller S, Franchet A, Hakkim A, Chen J, Drenkard E, Yu S, Schirmeier S, Li Z, Martins N, Ausubel FM, et al (2018a) Quorum-sensing regulator RhlR but not its autoinducer RhlI enables Pseudomonas to evade opsonization. EMBO Rep 19: e44880

Haller S, Franchet A, Hakkim A, Chen J, Drenkard E, Yu S, Schirmeier S, Li Z, Martins N, Ausubel FM, et al (2018b) Quorum-sensing regulator RhlR but not its autoinducer RhlI enables Pseudomonas to evade opsonization. EMBO Rep 19: e44880

Hamon Y, Trompier D, Ma Z, Venegas V, Pophillat M, Mignotte V, Zhou Z & Chimini G (2006) Cooperation between Engulfment Receptors: The Case of ABCA1 and MEGF10. PLOS ONE 1: e120

Hanson MA, Dostálová A, Ceroni C, Poidevin M, Kondo S & Lemaitre B (2019) Synergy and remarkable specificity of antimicrobial peptides in vivo using a systematic knockout approach. eLife 8: e44341

Hashimoto Y, Tabuchi Y, Sakurai K, Kutsuna M, Kurokawa K, Awasaki T, Sekimizu K, Nakanishi Y & Shiratsuchi A (2009) Identification of lipoteichoic acid as a ligand for draper in the phagocytosis of Staphylococcus aureus by Drosophila hemocytes. J Immunol 183: 7451–7460

Hedengren M, Asling B, Dushay MS, Ando I, Ekengren S, Wihlborg M & Hultmark D (1999) Relish, a central factor in the control of humoral but not cellular immunity in Drosophila. Mol Cell 4: 827–837

Hilu-Dadia R, Ghanem A, Vogelesang S, Ayoub M, Hakim-Mishnaevski K & Kurant E (2025) Santa-maria is a glial phagocytic receptor that acts with SIMU to recognize and engulf apoptotic neurons. Cell Rep 44: 115201

Honti V, Csordás G, Márkus R, Kurucz E, Jankovics F & Andó I (2010) Cell lineage tracing reveals the plasticity of the hemocyte lineages and of the hematopoietic compartments in Drosophila melanogaster. Mol Immunol 47: 1997–2004

Jumper J, Evans R, Pritzel A, Green T, Figurnov M, Ronneberger O, Tunyasuvunakool K, Bates R, Žídek A, Potapenko A, et al (2021) Highly accurate protein structure prediction with AlphaFold. Nature 596: 583–589

Jung S-H, Evans CJ, Uemura C & Banerjee U (2005) The Drosophila lymph gland as a developmental model of hematopoiesis. Development 132: 2521–2533

Kanampalliwar A, Shah M, Park Y, Jeong B, Lawson PA, Bell M, Fesko EM, Sainato A, Sainato D, Walker S, et al (2025a) The role of lipoteichoic acid in Staphylococcus aureus cell wall integrity. 2025.01.16.633316 doi:10.1101/2025.01.16.633316 [PREPRINT]

Kanampalliwar A, Shah M, Park Y, Jeong B, Lawson PA, Bell M, Fesko EM, Sainato A, Sainato D, Walker S, et al (2025b) The role of lipoteichoic acid in Staphylococcus aureus cell wall integrity. 2025.01.16.633316 doi:10.1101/2025.01.16.633316 [PREPRINT]

Kang JY, Nan X, Jin MS, Youn S-J, Ryu YH, Mah S, Han SH, Lee H, Paik S-G & Lee J-O (2009) Recognition of lipopeptide patterns by Toll-like receptor 2-Toll-like receptor 6 heterodimer. Immunity 31: 873–884

Kocks C, Cho JH, Nehme N, Ulvila J, Pearson AM, Meister M, Strom C, Conto SL, Hetru C, Stuart LM, et al (2005) Eater, a Transmembrane Protein Mediating Phagocytosis of Bacterial Pathogens in *Drosophila*. Cell 123: 335–346

Kristian SA, Datta V, Weidenmaier C, Kansal R, Fedtke I, Peschel A, Gallo RL & Nizet V (2005) d-Alanylation of Teichoic Acids Promotes Group A Streptococcus Antimicrobial Peptide Resistance, Neutrophil Survival, and Epithelial Cell Invasion. J Bacteriol 187: 6719–6725

Kurant E, Axelrod S, Leaman D & Gaul U (2008) The novel engulfment receptor Six-microns-under acts upstream of Draper in the glial phagocytosis of apoptotic neurons. Cell 133: 498–509

Kurucz E, Márkus R, Zsámboki J, Folkl-Medzihradszky K, Darula Z, Vilmos P, Udvardy A, Krausz I, Lukacsovich T, Gateff E, et al (2007) Nimrod, a putative phagocytosis receptor with EGF repeats in Drosophila plasmatocytes. Curr Biol CB 17: 649–654

Lagueux M, Perrodou E, Levashina EA, Capovilla M & Hoffmann JA (2000) Constitutive expression of a complement-like protein in toll and JAK gain-of-function mutants of Drosophila. Proc Natl Acad Sci U S A 97: 11427–11432

Lanot R, Zachary D, Holder F & Meister M (2001) Postembryonic hematopoiesis in Drosophila. Dev Biol 230: 243–257

Lee P-T, Zirin J, Kanca O, Lin W-W, Schulze KL, Li-Kroeger D, Tao R, Devereaux C, Hu Y, Chung V, et al (2018) A gene-specific T2A-GAL4 library for Drosophila. eLife 7: e35574

Lemaitre B, Nicolas E, Michaut L, Reichhart J-M & Hoffmann JA (1996) The Dorsoventral Regulatory Gene Cassette *spätzle/Toll/cactus* Controls the Potent Antifungal Response in Drosophila Adults. Cell 86: 973–983

Liegeois S & Ferrandon D (2022) Sensing microbial infections in the Drosophila melanogaster genetic model organism. Immunogenetics 74: 35–62

Logan MA, Hackett R, Doherty J, Sheehan A, Speese SD & Freeman MR (2012) Negative regulation of glial engulfment activity by Draper terminates glial responses to axon injury. Nat Neurosci 15: 722–730

MacDonald JM, Beach MG, Porpiglia E, Sheehan AE, Watts RJ & Freeman MR (2006) The Drosophila cell corpse engulfment receptor Draper mediates glial clearance of severed axons. Neuron 50: 869–881

Makhijani K, Alexander B, Tanaka T, Rulifson E & Brückner K (2011) The peripheral nervous system supports blood cell homing and survival in the Drosophila larva. Development 138: 5379–5391

Makhijani K & Brückner K (2012) Of blood cells and the nervous system: hematopoiesis in the Drosophila larva. Fly (Austin) 6: 254–260

Manaka J, Kuraishi T, Shiratsuchi A, Nakai Y, Higashida H, Henson P & Nakanishi Y (2004) Draper-mediated and phosphatidylserine-independent phagocytosis of apoptotic cells by Drosophila hemocytes/macrophages. J Biol Chem 279: 48466–48476

Mangahas PM & Zhou Z (2005) Clearance of apoptotic cells in Caenorhabditis elegans. Semin Cell Dev Biol 16: 295–306

Márkus R, Laurinyecz B, Kurucz E, Honti V, Bajusz I, Sipos B, Somogyi K, Kronhamn J, Hultmark D & Andó I (2009) Sessile hemocytes as a hematopoietic compartment in Drosophila melanogaster. Proc Natl Acad Sci U S A 106: 4805–4809

Matos RC, Nikolopoulos N, Perrier Q, Robert X, Gueguen-Chaignon V, Hirayama H, Leulier F, Grangeasse C, Guerardel Y & Ravaud S (2026) A DltE–DltD–DltX interaction network regulates lipoteichoic acid D-alanylation in Lactiplantibacillus plantarum and symbiotic drosophila growth promotion. 2026.03.19.713020 doi:10.64898/2026.03.19.713020 [PREPRINT]

Matskevich AA, Quintin J & Ferrandon D (2010) The Drosophila PRR GNBP3 assembles effector complexes involved in antifungal defenses independently of its Toll pathway activation function. Eur J Immunol 40: 1244–1254

McBride SM & Sonenshein AL (2011) The dlt operon confers resistance to cationic antimicrobial peptides in Clostridium difficile. Microbiology 157: 1457–1465

Melcarne C, Lemaitre B & Kurant E (2019a) Phagocytosis in *Drosophila*: From molecules and cellular machinery to physiology. Insect Biochem Mol Biol 109: 1–12

Melcarne C, Ramond E, Dudzic J, Bretscher AJ, Kurucz É, Andó I & Lemaitre B (2019b) Two Nimrod receptors, NimC1 and Eater, synergistically contribute to bacterial phagocytosis in Drosophila melanogaster. FEBS J 286: 2670–2691

Melcarne C, Ramond E, Dudzic J, Bretscher AJ, Kurucz É, Andó I & Lemaitre B (2019c) Two Nimrod receptors, NimC1 and Eater, synergistically contribute to bacterial phagocytosis in Drosophila melanogaster. FEBS J 286: 2670–2691

Melcarne C, Ramond E, Dudzic J, Bretscher AJ, Kurucz É, Andó I & Lemaitre B (2019d) Two Nimrod receptors, NimC1 and Eater, synergistically contribute to bacterial phagocytosis in Drosophila melanogaster. FEBS J 286: 2670–2691

Midega J, Blight J, Lombardo F, Povelones M, Kafatos F & Christophides GK (2013) Discovery and characterization of two Nimrod superfamily members in Anopheles gambiae. Pathog Glob Health 107: 463–474

Moeyaert B, Holt G, Madangopal R, Perez-Alvarez A, Fearey BC, Trojanowski NF, Ledderose J, Zolnik TA, Das A, Patel D, et al (2018) Improved methods for marking active neuron populations. Nat Commun 9: 4440

Moita LF, Wang-Sattler R, Michel K, Zimmermann T, Blandin S, Levashina EA & Kafatos FC (2005) In vivo identification of novel regulators and conserved pathways of phagocytosis in A. gambiae. Immunity 23: 65–73

Nagaosa K, Okada R, Nonaka S, Takeuchi K, Fujita Y, Miyasaka T, Manaka J, Ando I & Nakanishi Y (2011) Integrin βν-mediated Phagocytosis of Apoptotic Cells in Drosophila Embryos. J Biol Chem 286: 25770–25777

Needham AJ, Kibart M, Crossley H, Ingham PW & Foster SJ (2004) Drosophila melanogaster as a model host for Staphylococcus aureus infection. Microbiology 150: 2347–2355

Neuhaus FC & Baddiley J (2003) A continuum of anionic charge: structures and functions of D-alanyl-teichoic acids in gram-positive bacteria. Microbiol Mol Biol Rev MMBR 67: 686–723

Neyen C, Bretscher AJ, Binggeli O & Lemaitre B (2014a) Methods to study *Drosophila* immunity. Methods 68: 116–128

Neyen C, Bretscher AJ, Binggeli O & Lemaitre B (2014b) Methods to study Drosophila immunity. Methods 68: 116–128

Perron C, Carme P, Rosell AL, Minnaert E, Ruiz-Demoulin S, Szczkowski H, Neukomm LJ, Dura J-M & Boulanger A (2023) Chemokine-like Orion is involved in the transformation of glial cells into phagocytes in different developmental neuronal remodeling paradigms. Dev Camb Engl 150: dev201633

Petrignani B, Rommelaere S, Hakim-Mishnaevski K, Masson F, Ramond E, Hilu-Dadia R, Poidevin M, Kondo S, Kurant E & Lemaitre B (2021) A secreted factor NimrodB4 promotes the elimination of apoptotic corpses by phagocytes in Drosophila. EMBO Rep 22: e52262

Povelones M, Waterhouse RM, Kafatos FC & Christophides GK (2009) Leucine-rich repeat protein complex activates mosquito complement in defense against Plasmodium parasites. Science 324: 258–261

Ramond E, Petrignani B, Dudzic JP, Boquete J-P, Poidevin M, Kondo S & Lemaitre B (2020) The adipokine NimrodB5 regulates peripheral hematopoiesis in Drosophila. FEBS J 287: 3399–3426

Reed P, Veiga H, Jorge AM, Terrak M & Pinho MG (2011) Monofunctional transglycosylases are not essential for Staphylococcus aureus cell wall synthesis. J Bacteriol 193: 2549–2556

Romeo Y & Lemaitre B (2008) Drosophila immunity: methods for monitoring the activity of Toll and Imd signaling pathways. Methods Mol Biol 415: 379–394

Rommelaere S, Schüpfer F, Armand F, Hamelin R & Lemaitre B (2025) An updated proteomic analysis of Drosophila haemolymph after bacterial infection. Fly (Austin) 19: 2485685

Rosales C & Uribe-Querol E (2017) Phagocytosis: A Fundamental Process in Immunity. BioMed Res Int 2017: 9042851

Ryckebusch F, Tian Y, Rapin M, Schüpfer F, Hanson MA & Lemaitre B (2025) Layers of immunity: Deconstructing the Drosophila effector response. eLife 14

Sakr R, Monticelli S, Kizhakkenottiyath Shasthadevan S, Delaporte C, Zhang G, Tabiat T, Giangrande A & Cattenoz PB (2026) NimA promotes cell adhesion at the blood brain barrier of the Drosophila nervous system. EMBO Rep 27: 1648–1665

Schneewind O & Missiakas D (2014) Lipoteichoic Acids, Phosphate-Containing Polymers in the Envelope of Gram-Positive Bacteria. J Bacteriol 196: 1133–1142

Schröder NWJ, Morath S, Alexander C, Hamann L, Hartung T, Zähringer U, Göbel UB, Weber JR & Schumann RR (2003) Lipoteichoic acid (LTA) of Streptococcus pneumoniae and Staphylococcus aureus activates immune cells via Toll-like receptor (TLR)-2, lipopolysaccharide-binding protein (LBP), and CD14, whereas TLR-4 and MD-2 are not involved. J Biol Chem 278: 15587–15594

Shiratsuchi A, Mori T, Sakurai K, Nagaosa K, Sekimizu K, Lee BL & Nakanishi Y (2012) Independent Recognition of Staphylococcus aureus by Two Receptors for Phagocytosis in Drosophila. J Biol Chem 287: 21663–21672

Shklyar B, Levy-Adam F, Mishnaevski K & Kurant E (2013) Caspase Activity Is Required for Engulfment of Apoptotic Cells. Mol Cell Biol 33: 3191–3201

Shokal U & Eleftherianos I (2017) Evolution and Function of Thioester-Containing Proteins and the Complement System in the Innate Immune Response. Front Immunol 8

Somogyi K, Sipos B, Pénzes Z, Kurucz E, Zsámboki J, Hultmark D & Andó I (2008) Evolution of genes and repeats in the Nimrod superfamily. Mol Biol Evol 25: 2337–2347

Stephenson HN, Streeck R, Grüblinger F, Goosmann C & Herzig A (2022) Hemocytes are essential for Drosophila melanogaster post-embryonic development, independent of control of the microbiota. Dev Camb Engl 149: dev200286

Stroschein-Stevenson SL, Foley E, O’Farrell PH & Johnson AD (2006) Identification of Drosophila Gene Products Required for Phagocytosis of Candida albicans. PLoS Biol 4: e4

Troha K & Buchon N (2019) Methods for the study of innate immunity in Drosophila melanogaster. Wiley Interdiscip Rev Dev Biol 8: e344

Tung TT, Nagaosa K, Fujita Y, Kita A, Mori H, Okada R, Nonaka S & Nakanishi Y (2013) Phosphatidylserine recognition and induction of apoptotic cell clearance by Drosophila engulfment receptor Draper. J Biochem (Tokyo) 153: 483–491

Ulvila J, Vanha-Aho L-M & Rämet M (2011) Drosophila phagocytosis - still many unknowns under the surface. APMIS Acta Pathol Microbiol Immunol Scand 119: 651–662

Vlisidou I & Wood W (2015) Drosophila blood cells and their role in immune responses. FEBS J 282: 1368–1382

Westlake H, David F, Tian Y, Krakovic K, Dolgikh A, Juravlev L, Bournonville TE de, Carboni A, Melcarne C, Shan T, et al (2026) Reproducibility of Scientific Claims in Drosophila Immunity: A Retrospective Analysis of 400 Publications. eLife 15

Westlake H, Hanson MA & Lemaitre B (2024) The Drosophila immunity handbook EPFL Press

Winkelstein JA (1973) Opsonins: Their function, identity, and clinical significance. J Pediatr 82: 747–753

Wright AE & Douglas SR (1904) An experimental investigation of the rôle of the blood fluids in connection with phagocytosis. Proc R Soc Lond 72: 357–370

Zeigler DR, Prágai Z, Rodriguez S, Chevreux B, Muffler A, Albert T, Bai R, Wyss M & Perkins JB (2008) The Origins of 168, W23, and Other Bacillus subtilis Legacy Strains. J Bacteriol 190: 6983–6995

Zsámboki J, Csordás G, Honti V, Pintér L, Bajusz I, Galgóczy L, Andó I & Kurucz É (2013) Drosophila Nimrod proteins bind bacteria. Cent Eur J Biol 8: 633–645

