## Supplementary figures for "The NimB2 opsonin promotes *S. aureus* recognition by macrophages in *Drosophila melanogaster*"

**Appendix Figure S1**

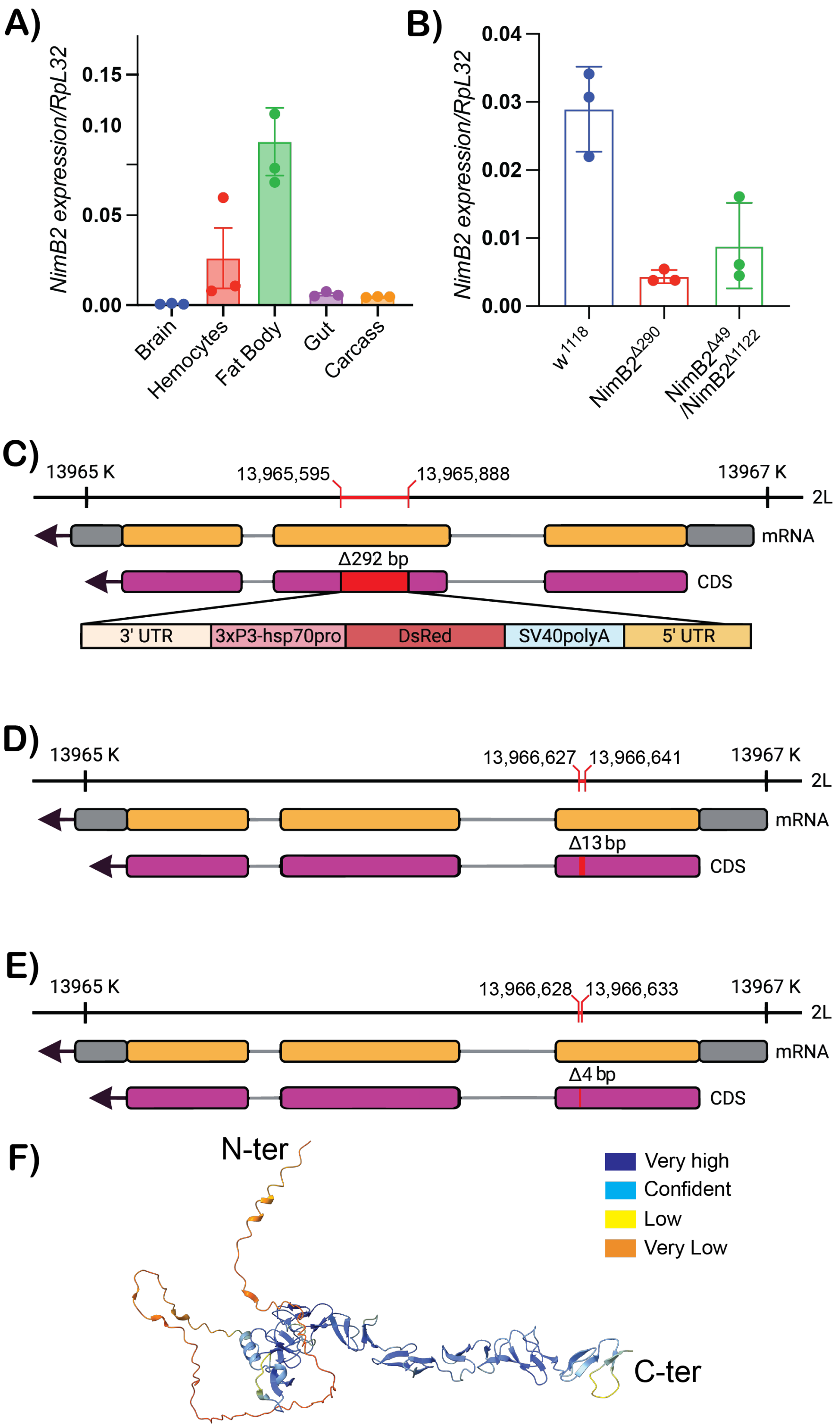


**Appendix Figure S1. Description of *NimB2* expression profile and *NimB2* mutant alleles.** (A) qPCR analysis of *NimB2* transcript levels in dissected tissues from third-instar larvae, confirming the enrichment of *NimB2* in the fat body, consistent with the *NimB2-Gal4* reporter expression pattern. (B) qPCR analysis of *NimB2* transcript levels in the indicated *NimB2* mutant alleles. (C) Schematic of the *NimB2* genomic locus depicting the CRISPR/Cas9-mediated *NimB2*$^{Δ290}$ mutation, in which a 292-bp segment of the *NimB2* coding sequence in exon 2 was replaced by a 3XP3-DsRed cassette. (D, E) Schematics of the *NimB2* genomic locus depicting the CRISPR/Cas9-mediated induced 13 bp and 4 bp deletion causing a frameshift in *NimB2*$^{Δ49}$ and *NimB2*$^{Δ1122}$ mutants, in exon 1 of the *NimB2* coding sequence. (F) AlphaFold prediction of NimB2 showing the predicted protein structure and per-residue confidence scores (pLDDT). Higher confidence is observed for the folded C-terminal region, whereas the N-terminal region shows lower confidence suggesting that it is disorder.

**Appendix Figure S2**

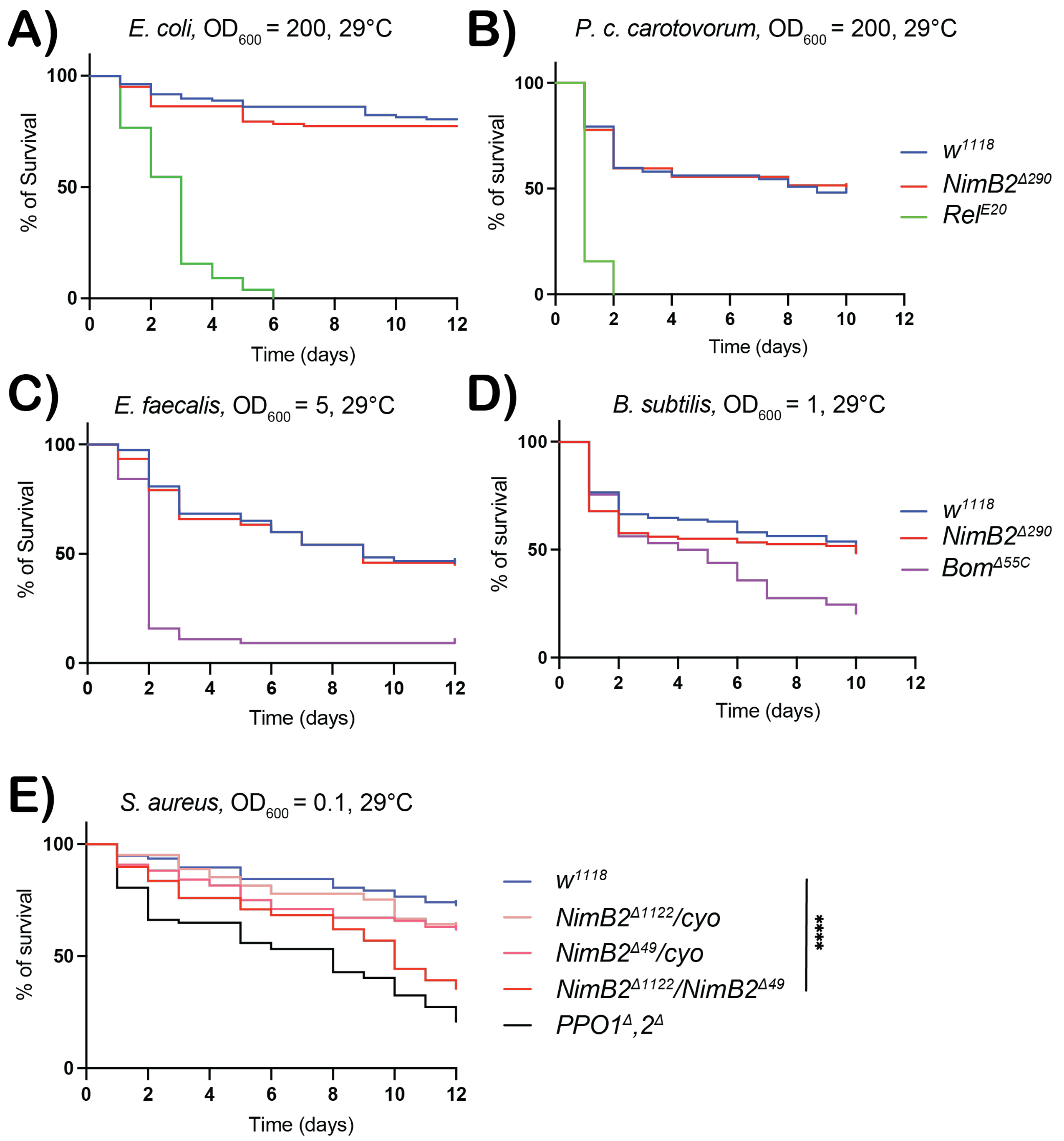


**Appendix Figure S2. *NimB2* mutants display a marked susceptibility specifically to *S. aureus*.** (A, B) *NimB2* mutants showed survival comparable to wild-type controls following infection with the Gram-negative bacteria, *Escherichia coli* and *P. c. carotovorum*. (C, D) *NimB2* mutants showed survival comparable to wild-type controls following infection with the Gram-positive bacteria, *Enterococcus faecalis* and *Bacillus subtilis*. (E) Trans heterozygous *NimB2$^{Δ49}$/NimB2$^{Δ1122}$* flies displayed an increased susceptibility to *S. aureus* compared with wild-type, or *NimB2$^{Δ1122}$/Cyo* and *NimB2$^{Δ49}$/Cyo* flies. This indicates a selective requirement for NimB2 in host defense against *S. aureus*. Survival distributions were compared with the *w$^{1118}$* wild-type control using the log-rank (Mantel–Cox) test. Optical density$_{600}$ of the bacterial pellet and temperature are indicated on the top of each panel. Statistical significance was defined as ns, not significant; P < 0.05 (*); P < 0.01 (**); P < 0.001 (***); and P < 0.0001 (****).

**Appendix Figure S3**

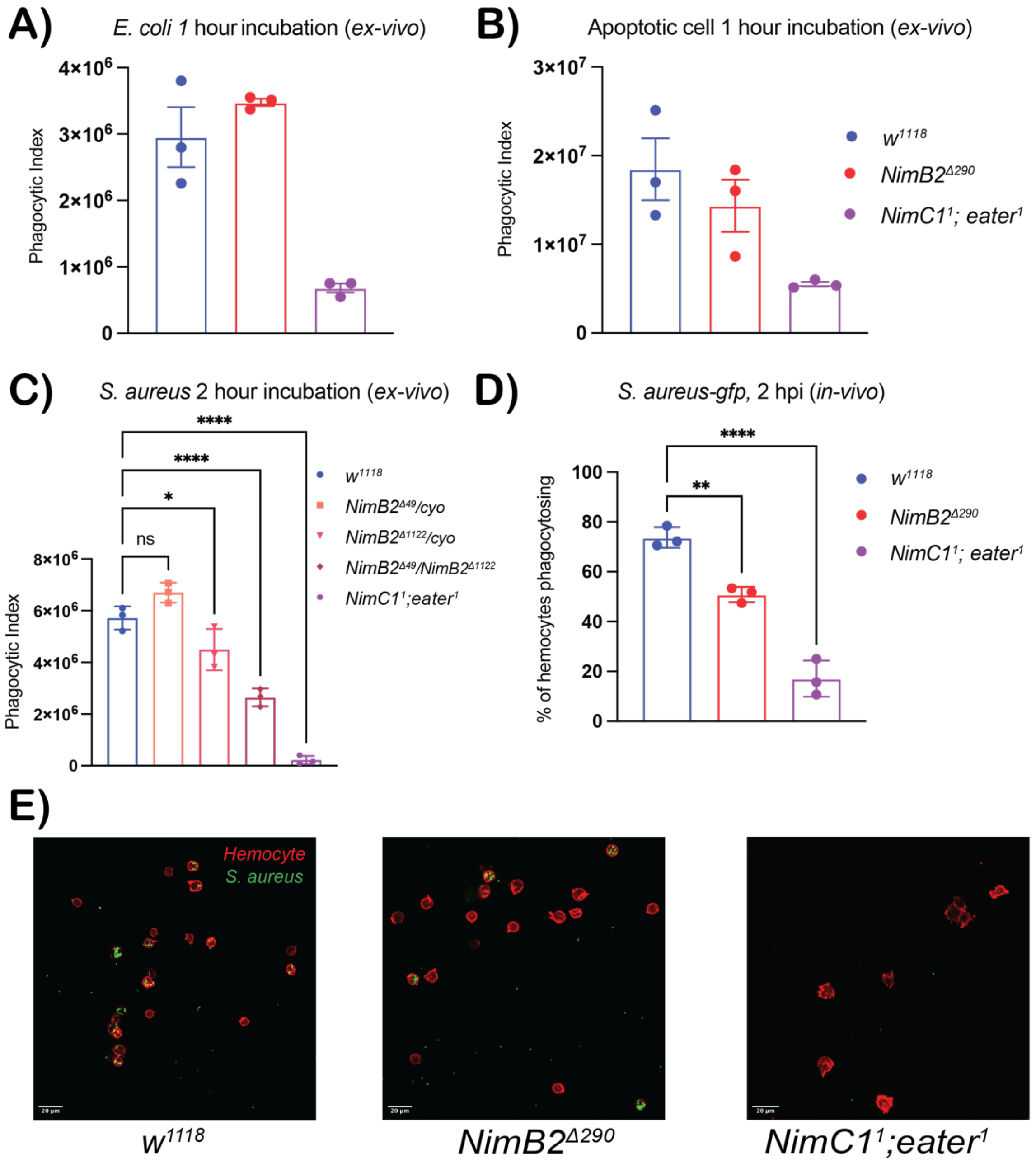


**Appendix Figure S3. NimB2 selectively promotes phagocytosis of *S. aureus* but not *E. coli* or apoptotic cells.** (A) *Ex vivo* phagocytosis assay using larval hemocytes from wild-type, NimB2$^{\Delta 290}$ and *NimC1$^{1}$, eater$^{1}$* larvae incubated with fluorescent *E. coli* bioparticles. The phagocytic index was not statistically different between *NimB2* mutant and the wild-type, indicating that NimB2 has no major role in *E. coli* phagocytosis. As expected, *NimC1; eater* double mutants display a marked defect in the engulfment of *E. coli* (Melcarne *et al*, 2019). (B*) Ex vivo* phagocytosis assay using apoptotic cell corpses with hemocytes from wild-type, NimB2$^{\Delta 290}$ and *NimC1$^{1}$, eater$^{1}$* third-instar larvae. In contrast to *NimC1$^{1}$, eater$^{1}$* mutations, the loss of *NimB2* did not affect the ability to phagocytose apoptotic corpses, indicating that NimB2 is not critical for efferocytosis. (C) *Ex vivo* phagocytosis assay using fluorescent *S. aureus* bioparticles, comparing wild-type and *NimB2$^{\Delta 49}$/NimB2$^{\Delta 1122}$* transheterozygous larvae, confirming the *S. aureus*-specific phagocytic defect in another *NimB2* deficient mutant background. (D, E) Quantification and representative images of *S. aureus-GFP* association with hemocytes using an *in vivo* assay phagocytic assay. Hemocytes from wild-type larvae show greater bacterial association than hemocytes *NimB2* mutants, while hemocytes from *NimC1, eater double-mutant larvae* showed almost bacterial association. In these experiments, we monitor only *S. aureus* GFP associated with hemocyte membrane because of the quenching of GFP in the acidic phagosome. Statistical significance was assessed by one-way ANOVA; ns, not significant. Statistical significance was defined as ns, not significant; $P < 0.05$ (*); $P < 0.01$ (**); $P < 0.001$ (***); and $P < 0.0001$ (****).

#### Appendix Figure S4

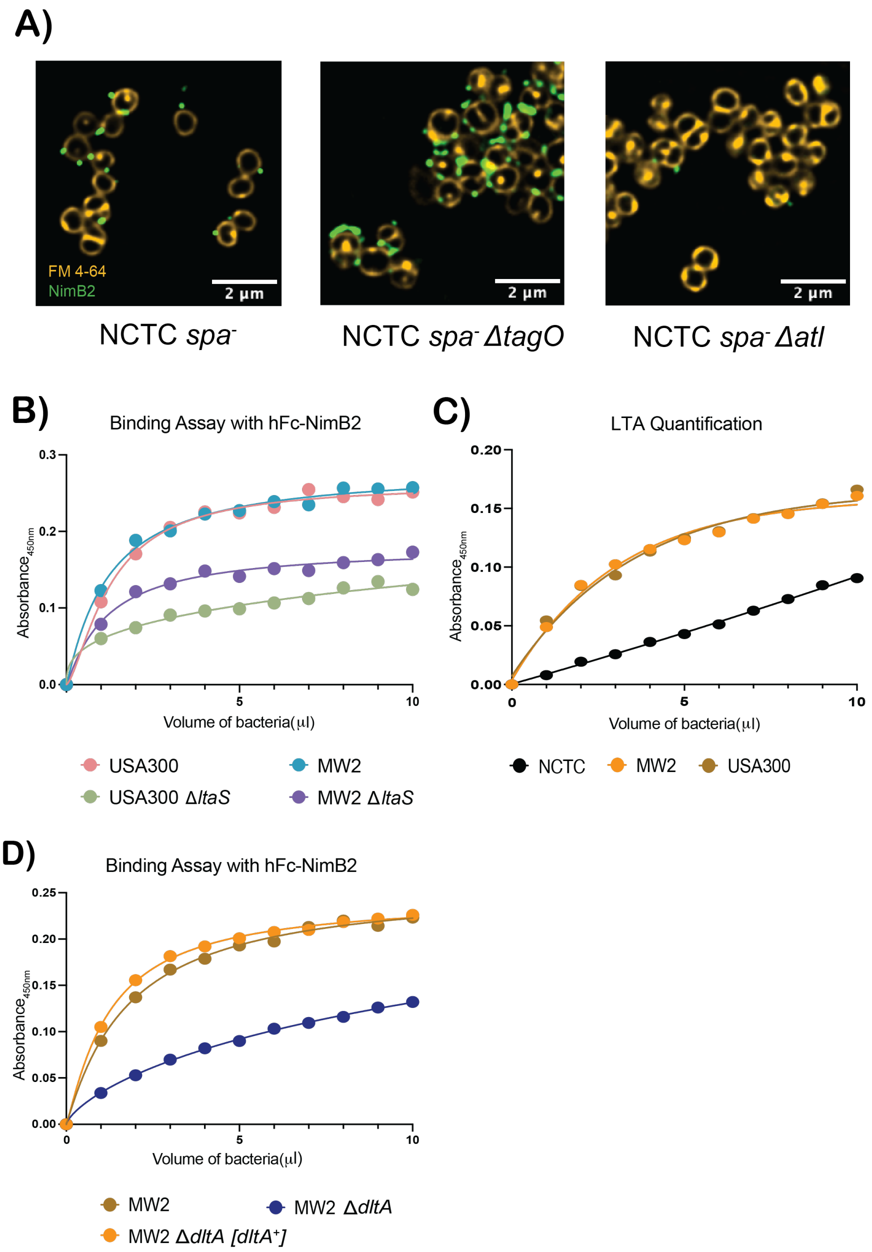


**Appendix Figure S4. Cell-wall determinants underlying NimB2 binding to *S. aureus*.** (A) Representative fluorescence microscopy images of NimB2 binding, to *spa*−, *spa*− Δ*tagO*, and *spa*− Δ*atl S. aureus* mutants labeled with FM4-64 (yellow). Loss of WTA increased accessibility of the NimB2-binding determinant, whereas loss of *atl* markedly reduced NimB2 binding. NimB2 was detected using a rabbit anti-NimB2 primary antibody and an Alexa Fluor 488 conjugated to secondary antibody anti-rabbit (green). Bacterial membrane was marked with FM 4 64 lipophilic dye. (B) ELISA-based binding assay showing NimB2 association with parental and Δ*ltaS S. aureus* strains in two genetic backgrounds. Loss of *ltaS* markedly reduced NimB2 binding compared with the corresponding parental strains. Binding curves represent pooled data from three independent experiments. C) Quantification of relative LTA abundance across three *S. aureus* strain backgrounds (NCTC, MW2, and USA300), showing higher LTA levels in USA300 and MW2 compared to NCTC. The differential LTA binding parallel the strain-dependent differences in NimB2 binding observed by Western blot (Fig. 7A) and ELISA assay (panel B) consistent with the notion that NimB2 binds to LTA dependent determinant. (D) ELISA-based binding assay showing NimB2 association with MW2 and MW2 Δ*dltA* strains as well as the complemented MW2 Δ*dltA [dltA*$^{+}$*]* strain. Loss of *dltA* reduced NimB2 binding, whereas complementation restored binding. Binding curves represent pooled data from three independent experiments. In ELISA assays shown in panels C and D, we used the hFc-NimB2 tagged version to reveal NimB2 binding to bacteria. The presence of Spa protein on MW2 and USA300 strains could explain the residual binding of NimB2 to Δ*ltaS and* Δ*dltA S. aureus.*

#### Appendix Figure S5

# A) *$w^{1118}$*

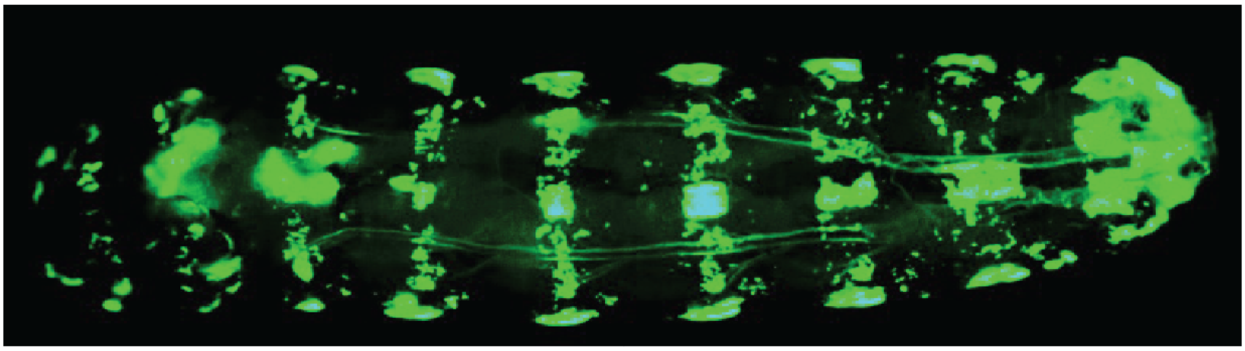

### B) *$NimB2^{\Delta 290}$*

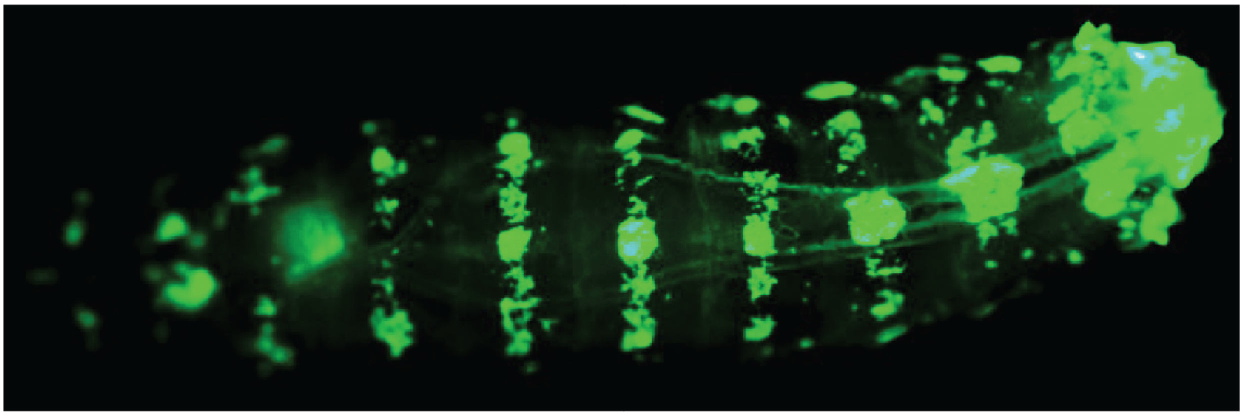

**Appendix Figure S5. Loss of *NimB2* does not affect hemocyte sessility in larvae.** Representative images of third-instar larvae showing hemocyte distribution revealed with the *Hml-Gal4, UAS-GFP* hemocyte marker. (A) Wild-type larvae and (B) *$NimB2^{\Delta 290}$* mutant larvae. In both genotypes, hemocytes display the characteristic sessile hemocyte pattern along the larval body wall, with no apparent alteration in hemocyte distribution in the absence of *NimB2*.
